# Resilient CD8^+^ T cells maintain a high cytotoxic capacity by balancing ROS via ME1 upregulation

**DOI:** 10.1101/2022.11.25.517988

**Authors:** Joanina K. Gicobi, Zhiming Mao, Grace DeFranco, Ying Li, Xin Liu, Jacob B. Hirdler, Vianca V. Vianzon, Emilia R. Dellacecca, Michelle A. Hsu, Whitney Barham, Yohan Kim, Feven Abraha, William S. Harmsen, Yiyi Yan, Roxana S. Dronca, Mojun Zhu, Svetomir N. Markovic, Aaron S. Mansfield, Yi Lin, Xiaosheng Wu, Dawn Owen, Michael P. Grams, Jacob J. Orme, Fabrice Lucien, Hu Zeng, Sean S. Park, Haidong Dong

**Affiliations:** Department of Immunology, Mayo Clinic College of Medicine and Science, Rochester, MN; Department of Urology, Mayo Clinic, Rochester, MN; Department of Biomedical Statistics and Informatics, Mayo Clinic, Rochester, MN; Department of Quantitative Health Sciences, Mayo Clinic, Rochester, MN; Division of Medical Oncology, Mayo Clinic, Jacksonville, FL; Division of Medical Oncology, Mayo Clinic, Rochester, MN; Department of Radiation Oncology, Mayo Clinic, Rochester, MN; Department of Rheumatology, Mayo Clinic, Rochester, MN

## Abstract

Cytotoxic T lymphocytes (CTL) are indispensable in anti-tumor immunity. Although CTLs are prone to exhaustion in patients with advanced cancer, T cell resiliency explains the presence of tumor-reactive CTLs that are less exhausted, capable of cytolytic function, expansion, and rebound in response to immunotherapy to reject metastatic malignances. However, the features of resilient T cells have not been clearly defined. In this report, we demonstrate that peripheral CX3CR1^+^ CD8^+^ T cells with low mitochondrial membrane potential rebounded CTL function quickly after radiation therapy in patients with large tumor burden portraying their functional resiliency. Furthermore, CX3CR1^+^ CD8^+^ T cell with low, but not high, mitochondrial membrane potential are highly cytotoxic, accumulate less reactive oxygen species (ROS), and express more Malic enzyme 1 (ME1). ME1 overexpression increases ATP production in a glycolysis-independent manner while concurrently curtailing excessive ROS in activated CD8^+^ T cells; and expands CX3CR1^+^NKG7^+^ effector CD8^+^ T cells with enhanced cytotoxicity. Importantly, transfection of *ME1* mRNA promotes tumoricidal activity in CD8^+^ T cells from patients with advanced cancers. Our study reveals a mechanism used by CTLs to balance excessive ROS via ME1 to maintain a metabolic and functional resiliency. Modification of ME1 expression in CTLs may be a novel method to improve the efficacy of cancer immunotherapy by preventing T cell exhaustion.

**Graphical abstract:** 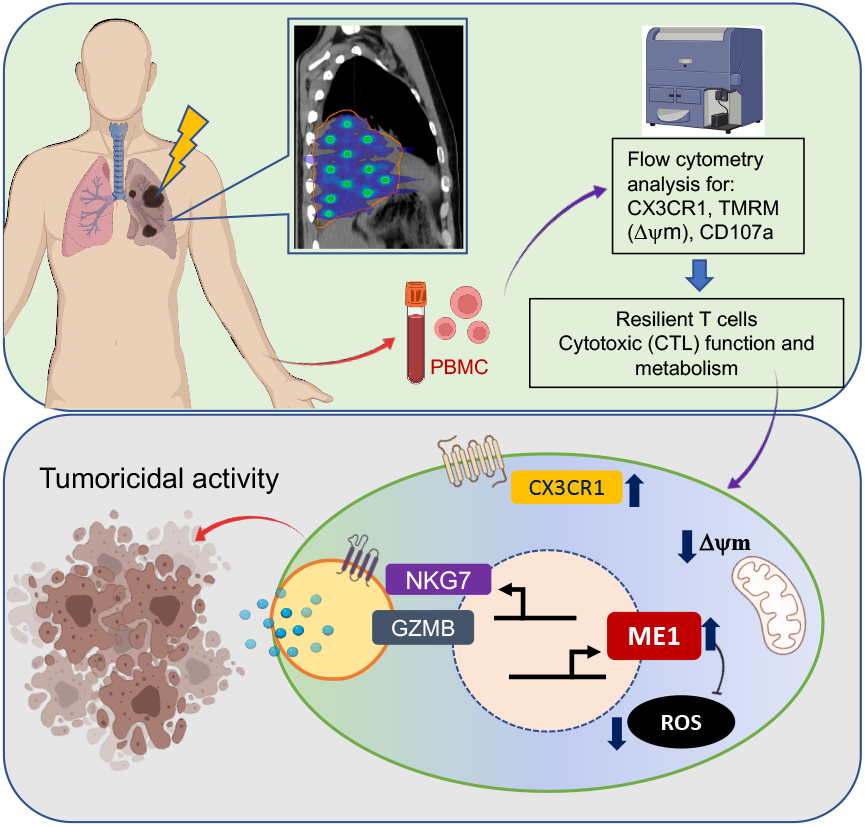

**Highlights:**

- CX3CR1^+^ and low Dy m identify functional resilient CD8^+^ T cells.
- Resilient CD8^+^ T cells are highly cytotoxic and have less ROS.
- Resilient CD8^+^ T cells express more ME1 that can balance extra ROS.
- ME1 overexpression can promote CTL function of CD8^+^ T cells.

## Introduction

Immunotherapies (immune checkpoint inhibitors (ICI), vaccines, or T cell therapies) are dependent on the presence of endogenous cytotoxic T lymphocytes (CTLs) to respond to ICI or to provide a substrate for adoptive cell transfer therapy^1^. Moreover, radiation therapy has been used frequently to reduce the tumor burden and to create opportunity for immunotherapy synergy^2^. To that end, the functionality of endogenous tumor-reactive CTLs plays a key role in predicting clinical response to combined radiation and immunotherapy^3,4^.

Recently, we have proposed the term “resilient T cells” (Trs) to describe a functional state of tumor-reactive cytotoxic T cells that are capable of responding to immunotherapy by withstanding tumor burden and enduring the harsh conditions of the tumor microenvironment ^5,6^. From there, a computational model of T cell resilience in multiple tumor types has been established to predict whether a cancer immunotherapy will be effective in certain patients ^7^. However, it is not clear whether Trs cells can be identified in the circulation of patients with large tumor burden (5-10 cm or more in diameter). Of note, the identification and characterization of circulating Trs cells in these patients is pivotal to clinical treatment plans because the presence of Trs cells is the foundation for the control of advanced cancer by combination immunotherapy.

It has been reported that successful ICI therapy expands tumor-reactive CD8^+^ T cells with an effector phenotype in the peripheral blood, which have the potential to replace exhausted T cells inside tumor tissue ^8–12^. To that end, we and others identified CX3CR1^+^ CD8^+^ T cells that are responsive to ICI therapy in both preclinical and clinical setting ^11,13,14^ Because CX3CR1^+^ CD8^+^ T cells are highly cytotoxic, proliferative, and active in preclinical models and in the peripheral blood of patients who respond to ICI therapy ^10,11,13^, we hypothesize that CX3CR1^+^ CD8^+^ T cells are prototypes of Trs cells in the circulation of patients with advanced cancers. A further characterization of CX3CR1^+^ CD8^+^ T cells will provide insights into the biology of Trs cells in circulation.

In this report, we found that post-radiation therapy in patients with advanced cancer, CX3CR1^+^ CD8^+^ T cells with low mitochondrial membrane potential (ΔΨm) undergo a rapid cytotoxic response. Upon further characterization of CD8^+^ T cells with low ΔΨm, we found that they are highly cytotoxic and accumulate less ROS (reactive oxygen species) but express more Malic enzyme 1 (ME1) compared to T cells with high ΔΨm. From there, we found that ME1 acts as an anti-oxidant in CD8^+^ T cells and expands CX3CR1^+^ NKG7^+^ effector CD8^+^ T cells with enhanced cytotoxicity. Importantly, ME1 overexpression introduced by *ME1* mRNA also increases mitochondrial-linked ATP while curtailing excessive ROS, thus enhancing the tumoricidal activity in CD8^+^ T cells from patients with advanced cancers. Our study suggests that resilient T cells balance excessive ROS upon antigen stimulation via upregulation of ME1 to maintain their metabolic fitness and retain a high cytotoxic capacity. Modification of ME1 expression in T cells may be a new method to avoid T cell exhaustion and improve the efficacy of cancer immunotherapy.

## Results

### CX3CR1 and low mitochondria membrane potential identify resilient CD8^+^ T cells in patients with advanced cancers

To reduce the tumor burden in patients with very large (5-10 cm or larger in diameter) or therapy-resistant tumors, patients were treated with a unique form of radiation therapy, spatially fractionated radiotherapy (SFRT) (**Fig. 1a**). SFRT is unique in that it intentionally delivers a heterogeneous dose distribution providing alternating regions of very high and low doses throughout the tumor^15^. SFRT can produce dramatic relief of severe symptoms, cause significant tumor regression, and achieve above average local control rates ^16^. The biologic mechanisms responsible for the effectiveness of SFRT are still unknown, but the heterogeneous dose distribution is hypothesized to be responsible for modifying tumor-reactive immune responses both locally and systemically.

**Fig. 1:**
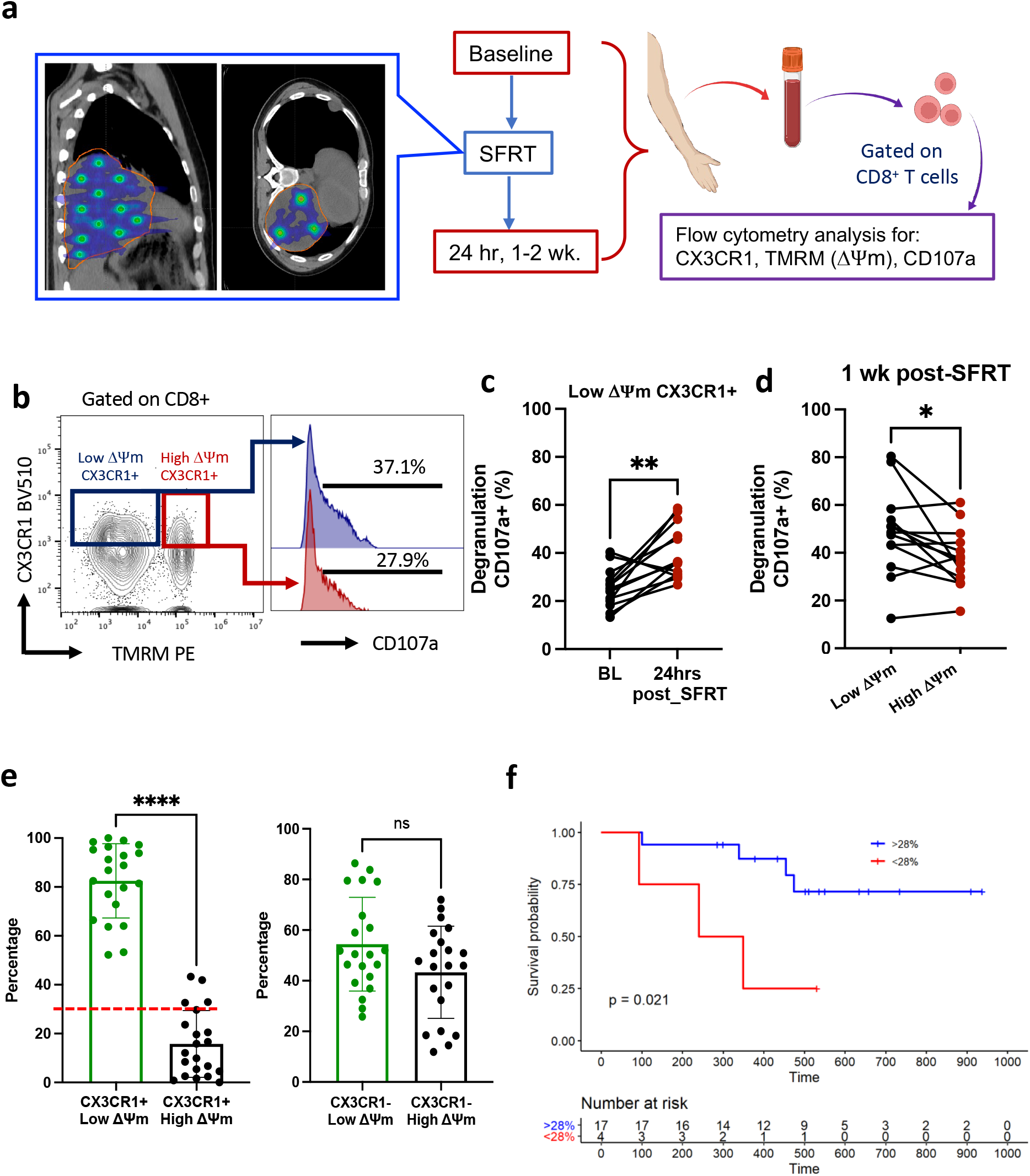
CX3CR1 and low ΔΨm identifies resilient CD8 T cells in patients with advanced cancers upon radiation therapy. **a**, A diagram of spatially fractionated radiotherapy (SFRT) therapy and study design for **b-c**. **b**, Degranulation assay (measured by CD107a expression) of CX3CR1^+^ CD8^+^ T cells with low or high ΔΨm. **c-d**, Summaries of percentages of CD107a^+^ cells among CX3CR1^+^CD8^+^ T cells with low or high ΔΨm from patients with advanced lung cancer and sarcoma (n=17) at baseline and 24 hours (hrs) post SFRT therapy (c), and one week (wk) post SFRT (d). Each line indicates one individual patient. **e**, Percentages of CX3CR1^+^ (left) and CX3CR1^-^ (right) CD8^+^ T cells with low or high ΔΨm from patients with melanoma receiving SBRT at baseline. **f**, Survival probabilities at baseline prior to stereotactic body radiotherapy (SBRT) in patients (n=20) with advanced melanoma. Kaplan-Meier estimates were used to estimate survival curves and a univariate Cox model was used to determine the association of patients and disease variables with the risk of each outcome of interest. Statistical significance was determined by Student’s Paired two-tailed t-test in **b-c**, and non-paired t-test in **d**, *P<0.05, **P<0.01, ****P<0.0001, ns: not significant.

We and others previously identified circulating CX3CR1^+^ CD8^+^ T cells that are less exhausted and demonstrate high cytotoxic capability in patients with advanced cancers such as melanoma and lung cancers ^10,11,13^. We hypothesized that a change in the functional state of CX3CR1^+^ CD8^+^ T cells may reflect an optimal response to a successful SFRT. Given the presence of large tumor burden in these patients in which compromised mitochondrial function has been linked to T cell exhaustion ^17–22^, we measured the CTL function of CX3CR1^+^ CD8^+^ T cells along with their mitochondrial membrane potential (ΔΨm) to determine their mitochondrial function. The CTL function of these T cells was measured by CD107a expression, which reflects degranulation and cytotoxicity. In lung cancer and sarcoma patients with large tumor burden, CX3CR1^+^ CD8^+^ T cells with low, but not high ΔΨm showed improved CTL function one day after SFRT (**Fig. 1a-c, Extended Data Fig. 1a**). Similarly, one week after SFRT we found CD8^+^ T cells with low ΔΨm showed improved CTL function compared to cells with high ΔΨm (**Fig. 1d**). In another cohort of patients with advanced melanoma whom were resistant to ICI therapy, we found that CX3CR1^+^ CD8^+^ T cells, but not CX3CR1^-^ CD8^+^ T cells, exhibit a low ΔΨm phenotype in the peripheral blood (**Fig. 1d**) prior to radiation therapy. We treated this cohort of patients with stereotactic body radiotherapy (SBRT) to reduce tumor burden. From there, we found patients with a higher frequency (>28%, cut-off line indicated in **Fig. 1e**) of CX3CR1^+^ CD8^+^ T cells with low ΔΨm at baseline demonstrated superior overall survival (OS) in response to SBRT (**Fig. 1f**). On the other hand, we found a gradually decreasing frequency of CX3CR1^+^ CD8^+^ T cells with low ΔΨm in responders to ICI therapy as the tumor burden reduced, but fluctuating frequency in non-responders and persistence of tumor burden (**Extended Data Fig. 1b-c**). Taken together our clinical observations suggest that CX3CR1^+^ CD8^+^ T cells with low ΔΨm are present in patients with large tumor burdens, and demonstrate a resiliency in their cytotoxic function after radiation therapy.

### Resilient CD8^+^ T cells are highly cytotoxic and less exhausted

Although murine CD8^+^ T cells with low ΔΨm demonstrates a higher antitumor activity ^23^, the cytotoxic functionality of human CD8^+^ T cells with low or high ΔΨm has not been clearly defined. To understand whether the low ΔΨm feature of CD8^+^ T cells is associated with their functional state, we sorted CD8^+^ T cells isolated from healthy donors into low or high ΔΨm using TMRM staining as reported ^23^ (**Fig. 2a; Extended Data Fig. 2a**). First, we performed bulk RNA-seq analysis to compare the transcriptional differences between CD8^+^ T cells with low and high ΔYm. We found CD8^+^ T cells with low ΔΨm expressed more genes that code for cytotoxic effector molecules such as *GZMB, PRF1*, and *NKG7* than CD8^+^ T cells with high ΔΨm (**Fig. 2b**). We then confirmed a higher expression of granzyme B in CD8^+^ T cells with low ΔΨm compared to CD8^+^ T cells with high ΔΨm using RT-PCR and flow cytometry (**Fig. 2c-d**). In a T cell-mediated tumor cytotoxicity assay following a brief T cell activation with anti-CD3/CD28, we found that CD8^+^ T cells with low ΔΨm demonstrated 1.5-fold higher cytolytic activity compared to CD8^+^ T cells with high ΔΨm in the killing of two tumor cell lines (breast cancer and prostate cancer) (**Fig. 2e**). Also, we observed significantly increased degranulation of CD8^+^ T cells with low ΔΨm compared to CD8^+^ T cells with high ΔΨm upon T cell activation (**Fig. 2f**). In addition, gene set enrichment analysis (GSEA) showed CD8^+^ T cells with low ΔΨm are enriched for mediators of inflammatory responses including Interferon (IFN) and IL-2/STAT5 pathways (**Fig. 2g; Extended Data Fig. 2b**). We determined the phenotype of human CD8^+^ T cells with low or high ΔΨm and found that CD8^+^ T cells with low ΔΨm cells were enriched in both Temra-like (CCR7^-^CD45RA^+^) and Tnaive-like (CCR7^+^CD45RA^+^) subsets of CD8^+^ T cells, indicating a limitation of using CCR7 and CD45RA to identify a functional state of resilient T cells (**Extended Data Fig. 2c**). Overall, our results suggest that human CD8^+^ T cells with low ΔΨm are highly cytotoxic and more responsive to inflammatory signals.

**Fig. 2:**
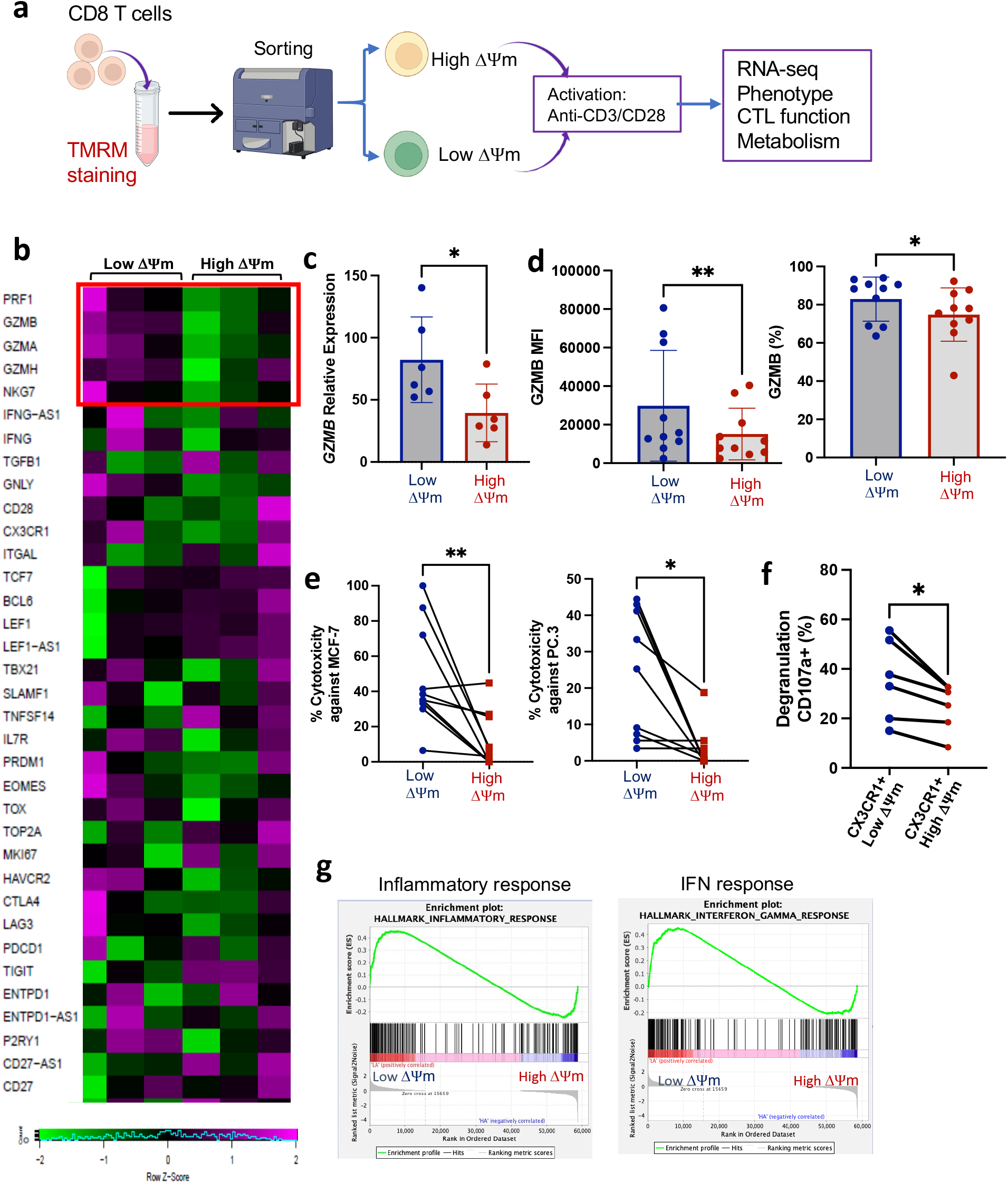
Resilient CD8 T cells have high cytolytic capability. **a**, Experimental schematic. **b**, Heatmap of differentially expressed genes from bulk RNA-sequencing (RNA-seq) of anti-CD3/CD28 activated CD8+ T cells sorted according to low and high ΔΨm (N=3). Framed area indicates genes that encode cytotoxic molecules**. c**, Relative GZMB mRNA expression in 5 donors. **d**, Expression of granzyme B protein in activated CD8^+^ T cells with low or high ΔΨm as analyzed by flow cytometry for MFI (median fluorescent intensity) and percentage of positive cells. **e**, Cytotoxicity assay of CD8^+^ T cells with low or high ΔΨm against tumor cells (MCF-7 breast and PC.3 prostate cancer) at a ratio of 1:20 (tumor:effector) in a 4-hour Calcein release assay (n=9-10). **f**, Degranulation assay of CX3CR1^+^ CD8^+^ T cells with low or high ΔΨm as measured by CD107a expression. **g**, Hallmark Gene Set Enrichment Analysis (GSEA) of RNA-seq data from three healthy donors using IPA analysis software. Statistical significance was determined by Student’s paired (**f**) or non-paired (**c-e**) two-tailed t-test, *P<0.05, **P<0.01.

Given the high cytotoxic capability of CD8^+^ T cells with low ΔΨm, we investigated whether they are prone to exhaustion upon activation. Interestingly, CD8^+^ T cells with low ΔΨm had lower expression of exhaustion markers such as TOX, a causative factor of T cell exhaustion^24^, and PD-1 (**Fig. 3a, b**) compared to cells with high ΔΨm. Interestingly, we found that CD8^+^ T cells with low ΔΨm are enriched with a TCF-1^+^PD-1^+^ stem-like phenotype that has been reported to be responsive to immunotherapy (**Fig. 3c**) ^23^. To confirm this possibility, we compared the cytotoxicity of CD8^+^ T cells with low or high ΔΨm in a co-culture with anti-PD-1 or anti-PD-L1 antibody. We found the cytotoxicity of CD8^+^ T cells with low ΔΨm increased in the presence of anti-PD-L1 or anti-PD-1 antibody compared to CD8^+^ T cells with high ΔΨm, although only the anti-PD-L1 group reached a statistical significance (**Fig. 3d**). Additionally, a pathway analysis of our RNA-seq data revealed an enrichment for the anti-PD-1/L1 cancer immunotherapy pathway in CD8^+^ T cells with low ΔΨm cells compared to high ΔΨm cells (**Extended Data Fig. 3a**). Since the transcription factors Eomesodermin (Eomes) and T-bet regulate the expression of effector genes in CD8^+^ T cells ^25–28^ and the ratio of Eomes/T-bet may determine the state of T cell exhaustion and immune pathology ^29,30^, we measured and compared Eomes and T-bet expression between CD8^+^ T cells with low or high ΔΨm. Although Eomes and T-bet expression at an mRNA level were significantly higher in CD8^+^ T cells with low ΔΨm compared to high ΔΨm cells, the protein levels of Eomes and T-bet were comparable between CD8^+^ T cells with low or high ΔΨm (**Fig. 3e-f**). These data suggest that although CD8^+^ T cells with low ΔΨm seem to be programmed with a high capability of effector function that could lead to exhaustion (based on high Eomes transcription), their functional state may be regulated by alternative mechanisms preventing them from entering exhaustion.

**Fig. 3:**
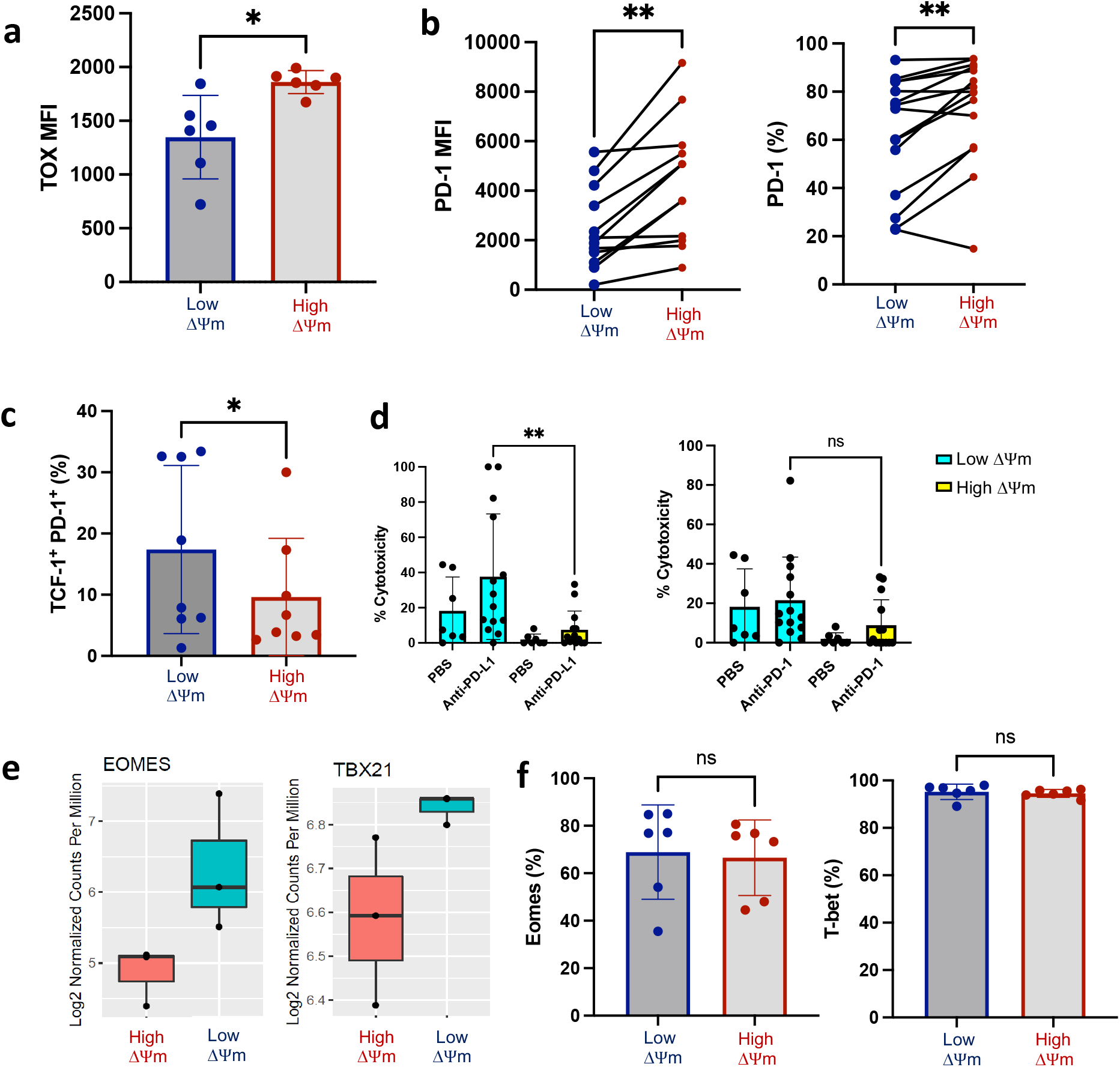
Resilient CD8 T cells are not prone to be exhausted. **a-b**, TOX (a) and PD-1 (b) expression by activated CD8^+^ T cells with low and high ΔΨm as analyzed by flow cytometry in median fluorescent intensity (MFI) and frequency (%). **c**, Frequency of TCF-1^+^ PD-1^+^ among CD8^+^ T cells with low or high ΔΨm at activated state. **d**, CD8^+^ T cells with low or high ΔΨm were activated with anti-CD3/CD28 beads in the presence of anti-PD-1 (Pembro) or anti-PD-L1 (Atezo) for 72 hours followed by inclusion in a cytotoxicity assay at a ratio of 1:20 (tumor:effector) against PC.3 tumor cells (n=7-13). **e-f**, Box plots of bulk RNA-sequencing transcripts (**e**) and flow cytometry analysis (**f**) of Eomes and T-bet in CD8^+^ T cells with low or high ΔΨm at activated state (n=6). Statistical significance was determined by Student’s Paired (**a**) and non-paired (**b-d, f**) two-tailed t-test, *P<0.05, **P<0.01, ns: not significant.

### Resilient CD8^+^ T cells have lower glycolysis and less ROS

As the levels of ΔΨm of CD8^+^ T cells reflect the metabolic state of T cells^23,31^, we compared the usage of glycolysis, measured by extracellular acidification rates (ECAR), and mitochondrial respiration, measured by the oxygen consumption rates (OCR), between CD8^+^ T cells with low ΔΨm or high ΔΨm. Effector CD8^+^ T cells rapidly utilize glycolysis as a quick, but inefficient, means of ATP production to carry out their effector functions^32–34^. However, a recent report argues that activated human CD8^+^ T cells increase their metabolic pathways in both glycolysis and oxidative phosphorylation ^35^. We found CD8^+^ T cells with low ΔΨm had lower glycolysis and glycolytic capacity than CD8^+^ T cells with high ΔΨm (**Fig. 4a, b**), while their OCR was comparable between CD8^+^ T cells with low ΔΨm or high ΔΨm (**Fig. 4c, d**). However, the OCR/ECAR ratio was higher in CD8^+^ T cells with low ΔΨm compared to high ΔΨm cells (**Fig. 4f**), implying that the resilient CD8^+^ T cells may utilize more mitochondrial respiration for energy needs instead of glycolysis. Accordingly, mitochondrial-linked ATP and spare respiratory capacity were comparable between CD8^+^ T cells with low ΔΨm and high ΔΨm (**Fig. 4d, f**). Given the high cytotoxic capacity of CD8^+^ T cells with low ΔΨm, it is unexpected that they would exhibit low glycolysis. Since GLUT1 levels were comparable between CD8^+^ T cells with low ΔΨm and high ΔΨm (**Fig. 4g**), it is possible that the lower glycolysis of CD8^+^ T cells with low ΔΨm is not due to a lower intake of glucose, but rather a preference for lower lactate production via glycolysis which could result in Warburg effects that limit their functionality^36^. Our data suggests that CD8^+^ T cells with low ΔΨm do not have increased glycolysis, but rather meet their energy requirements via mitochondrial-linked respiration.

**Fig. 4:**
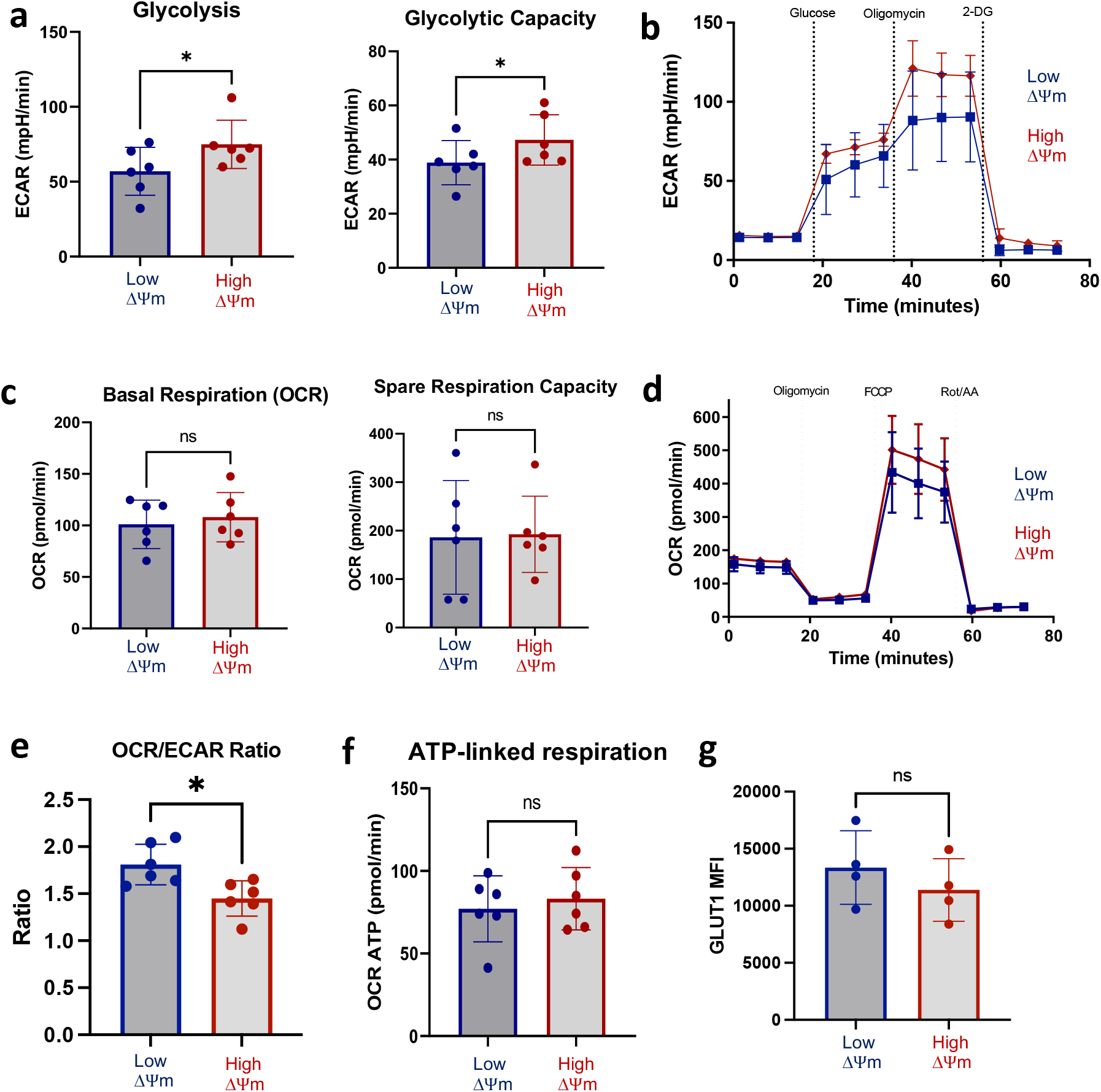
Resilient CD8^+^ T cells have lower glycolytic capacity. **a-b**, Glycolysis, and glycolytic capacity of sorted CD8^+^ T cells with low and high ΔΨm after activation with anti-CD3/CD28 beads as measured with extracellular acidification rate (ECAR) analysis and shown with a representative of Seahorse graph (**b**, n=6). **c**, Basal respiration, and spare respiratory capacity of sorted CD8^+^ T cells with low and high ΔΨm after activation as measured with Mitochondrial Oxygen Consumption Rate (OCR) analysis. **d**, Representative graph of OCR as measured by MitoStress Test (n=6). **e**, OCR/ECAR ratio calculated from the basal respiration values (n=6). **f**, Mitochondrial ATP of activated CD8^+^ T cells obtained from OCR analysis (n=6). **g**, GLUT1 expression (M (median)FI) in activated CD8^+^ T cells with low and high ΔΨm (n=4). Statistical significance was determined by Student’s paired two-tailed t-test; *P<0.05, ns: not significant.

Consistent with our observations above, our GSEA analysis shows that the ROS pathway was enriched in CD8^+^ T cells with high ΔΨm compared to those with low ΔΨm (**Fig. 5a**). To confirm this finding, we directly measured ROS levels in CD8^+^ T cells with low ΔΨm or high ΔΨm. We found both cytosolic ROS and mitochondrial ROS to be lower in resting and activated CD8^+^ T cells with low ΔΨm compared to CD8^+^ T cells with high ΔΨm (**Fig. 5b-c**), and that these lower levels of ROS were maintained for up to 7 days during the *in vitro* culture (**Fig. 5c**). To that end, we measured and compared the mass and numbers of mitochondria between CD8^+^ T cells with low and high ΔΨm. Although there was a trend for decreased mitochondrial mass and numbers of mitochondria in CD8^+^ T cells with low ΔΨm (**Fig. 5d-e**), this difference did not reach statistical significance, suggesting other mechanisms, beyond mitochondrial volume, may contribute to the lower levels of ROS.

**Fig. 5:**
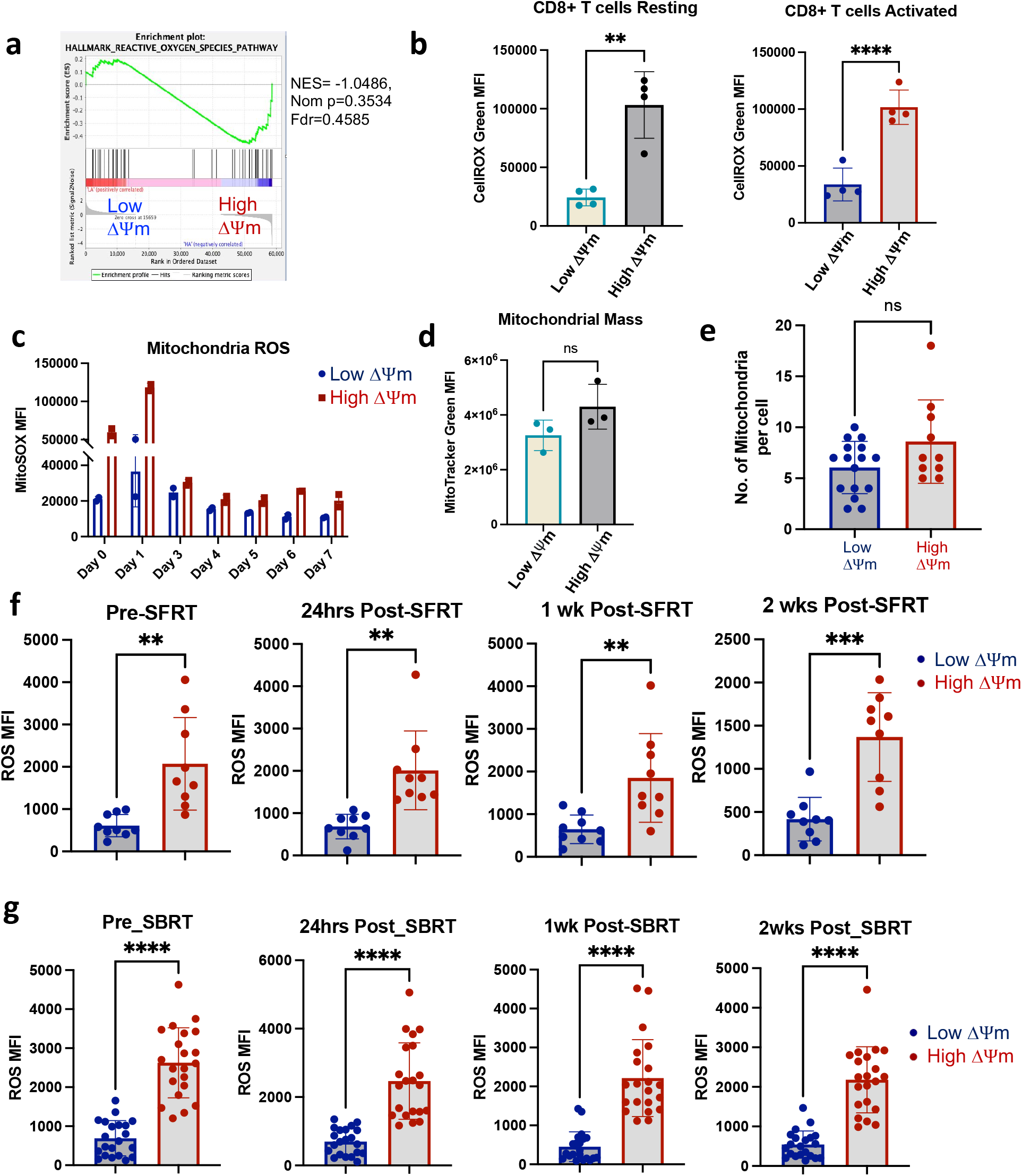
Resilient CD8 T cells have lower cytosolic and mitochondrial ROS. **a**, Hallmark GSEA analysis of reactive oxygen species (ROS) pathway genes from bulk RNA-sequencing data. **b**, Cytosolic ROS was measured by flow cytometry using CellROX Green and shown as MFI in both resting and activated CD8^+^ T cells. **c**, Mitochondrial ROS was measured by flow cytometry using MitoSOX over one week in culture media. **d**, Mitochondrial mass was measured using MitoTracker Green with flow cytometry and shown as M (median)FI. **e**, Number of mitochondria per cell was counted via Transmission electron microscopy (TEM) among 10-16 view fields. **f-g**, Cytosolic ROS in CD8^+^ T cells with low or high ΔΨm were measured from the PBMCs isolated from patients with lung cancer or sarcoma receiving spatially fractionated radiotherapy (SFRT) (**f**, n=9), and from patients with prostate cancer receiving stereotactic body radiotherapy (SBRT) (**g**, n=21). Statistical significance was determined with Student’s Nonpaired two-tailed t-test; **P<0.01, ***P<0.001, ****P<0.0001, ns: not significant.

Since excessive ROS signals result in DNA damage and functional exhaustion in T cells ^22,37,38^, the ability to maintain a lower ROS level, in the context of a higher OCR/ECAR ratio, could be a unique feature of resilient CD8^+^ T cells with low ΔΨm. Harsh tumor microenvironments lead to increased ROS in T cells, but CD8^+^ T cells with low ΔΨm may be enabled to tolerate these conditions. To test whether this feature is retained in T cells from patients with large tumor burdens, we measured and compared the levels of ROS in CD8^+^ T cells with low or high ΔΨm isolated from patients with advanced lung cancers and sarcoma before and after SFRT. Interestingly, a lower level of ROS was consistently maintained in CD8^+^ T cells with low ΔΨm compared to cells with high ΔΨm before and after SFRT (**Fig. 5f**, n=9). Similar findings were confirmed in another cohort of patients with advanced prostate cancer receiving SBRT (**Fig. 5f**, n=21). Taken together, our results suggest that CD8^+^ T cells with low ΔΨm selectively use mitochondrial respiration to meet their energy needs but have a capability to balance ROS levels to avoid an excessive ROS accumulation.

### ME1 is upregulated in Resilient CD8^+^ T cells

To determine how CD8^+^ T cells with low ΔΨm can maintain their ATP production via mitochondrial respiration, while also keeping a lower ROS level, we performed RNA-seq analysis and found ME1 to be among the most upregulated genes in the CD8^+^ T cells with low ΔΨm cells in both resting and activated states compared to CD8^+^ T cells with high ΔΨm (**Fig. 6a-b**). Since ME1 alone has the potential to regulate metabolism and ROS among these upregulated genes^39,40^, we focused on ME1 and confirmed its higher expression in CD8^+^ T cells with low ΔΨm cells compared to high ΔΨm cells via quantitative RT-PCR and Western blot in both resting and activated CD8^+^ T cells (**Fig. 6c-d**).

**Fig. 6:**
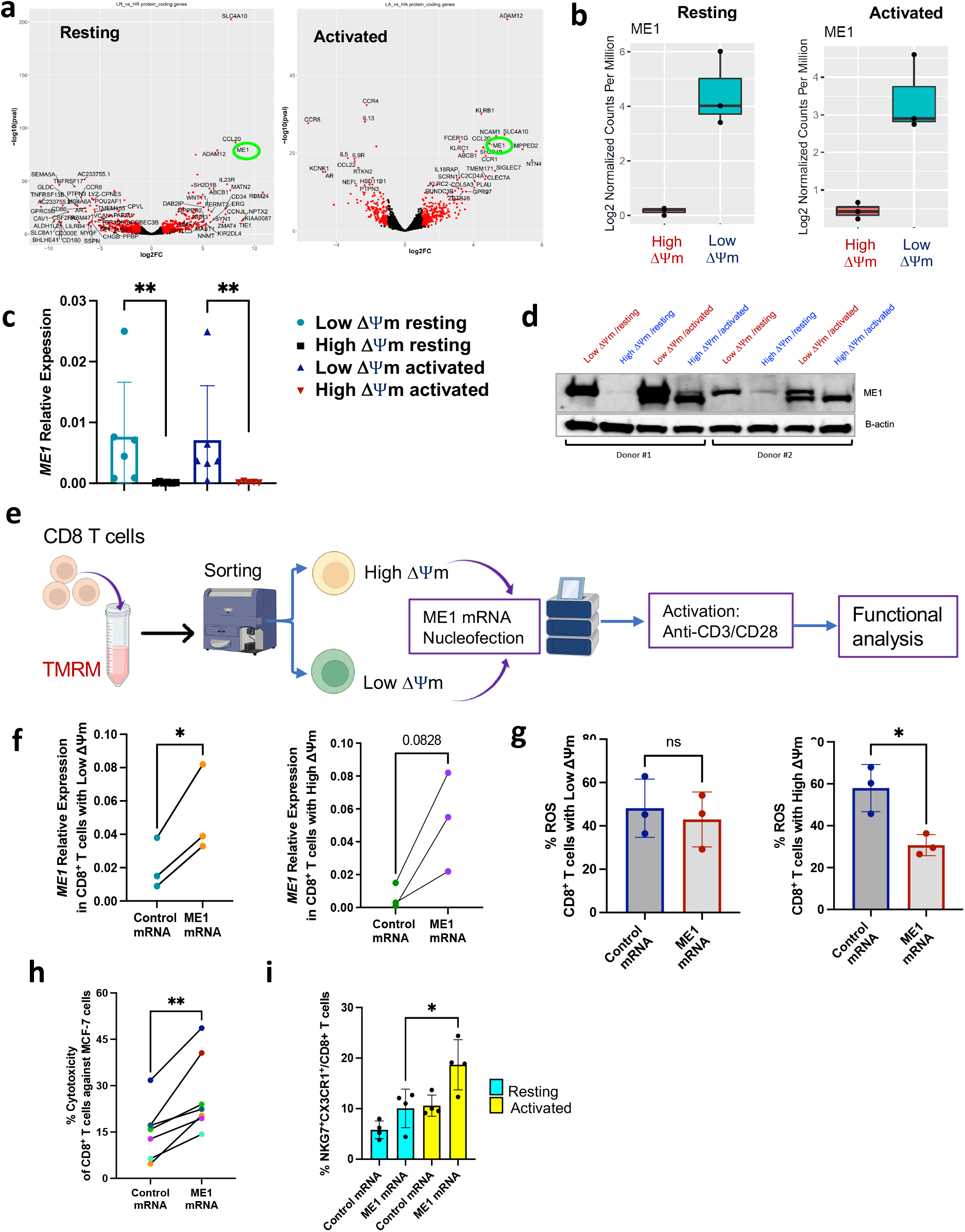
Resilient CD8 T cells express more ME1 to regulate ROS and promote CTL function. **a**, Volcano plots of bulk RNA-sequencing (RNA-seq) of CD8^+^ T cells with low or high ΔΨm at resting and activated state (n=3) (ME1 in green circle). **b**, Box plots of ME1 expression analysis with bulk RNA-seq as in (**a**). **c**, Relative ME1 mRNA expression measured with qRT-PCR. **d**, Western blot of ME1 expression as in (**c**) in two donors. **e**, Schematic of functional study of T cells with overexpression of ME1. **f**, ME1 overexpression measured by qRT-PCR in low and high ΔΨm cells. **g, Reactive oxygen species (**ROS) levels (%) measured in CD8^+^ T cells with low or high ΔΨm isolated from healthy donors (n=3) after transfection with control or ME1 mRNA. **h**, Cytotoxicity assay of CD8^+^ T cells isolated from healthy donors (n=6) after transfection with control or ME1 mRNA in killing tumor cells (MCF-7) at a ratio of 1:10 of tumor:effector cells. **i**, % CX3CR1^+^ NKG7^+^ among CD8^+^ T cells after transfection with control or ME1 mRNA as measured by flow cytometry (n=4). Statistical significance was determined by Student’s Paired two-tailed t-test; *P<0.05, **P<0.01, ns: not significant.

To examine the consequence of ME1 expression in CD8^+^ T cells with low or high ΔΨm, we sorted CD8^+^ T cells with low or high ΔΨm from healthy donors followed with ME1 mRNA transfection and functional analysis (**Fig. 6e**). The overexpression of ME1 was confirmed by RT-PCR (**Fig. 6f**). Interestingly, we found that ME1 overexpression reduced ROS levels only in CD8^+^ T cells with high ΔΨm, but not in CD8^+^ T cells with low ΔΨm (**Fig. 6g**). Of note, although ME1 has the capability to reduce ROS levels in CD8^+^ T cells, ME1 did not completely abolish ROS there, suggesting ME1 is a regulator of ROS rather than a scavenger of ROS in T cells. Since a basal level of ROS is also required for T cell activation^41^, we determined whether ME1 overexpression would affect the functionality of CD8^+^ T cells. To do so, we measured and compared the T cell-mediated cytotoxicity between CD8^+^ T cells transfected with control or ME1 mRNA. The overexpression of ME1 in CD8^+^ was confirmed via western blot (**Extended Data Fig. 4a-b**). Interestingly, we found that ME1 overexpression significantly increased the cytotoxicity of CD8^+^ T cells against tumor cells *in vitro* (**Fig. 6h**). Accordingly, we found ME1 overexpression significantly increased the frequency of highly cytotoxic CX3CR1^+^ NKG7^+^ CD8^+^ T cells ^42^ upon T cell activation (**Fig. 6i**). Our results suggest that ME1 may not only regulate the level of ROS in CD8^+^ T cells according to their mitochondrial potential, but also enhance CTL functionality.

### ME1 increases metabolic output of CD8^+^ T cells without increasing ROS

To understand how ME1 enhances CTL function of CD8^+^ T cells, we examined whether ME1 regulates T cell metabolism upon activation, being that ME1 is a cytosolic enzyme that converts malate into pyruvate and produces NAPDH^39,40^. Interestingly, we found that CD8^+^ T cells with ME1 overexpression had higher basal and maximal respiration, as well as a higher ATP-linked respiration, but their spare respiratory capacity did not increase significantly (**Fig. 7a; Extended Data Fig. 4c**). On the other hand, we found that ME1 overexpression did not affect the glycolysis and glycolytic capacity in CD8^+^ T cells (**Fig. 7b**). Importantly, the increased mitochondrial respiration induced by ME1 overexpression did not result in an elevated level of cytosolic ROS among total CD8^+^ T cells at either the activation or resting stage (**Fig. 7c; Extended Data Fig. 4d**).

**Fig. 7:**
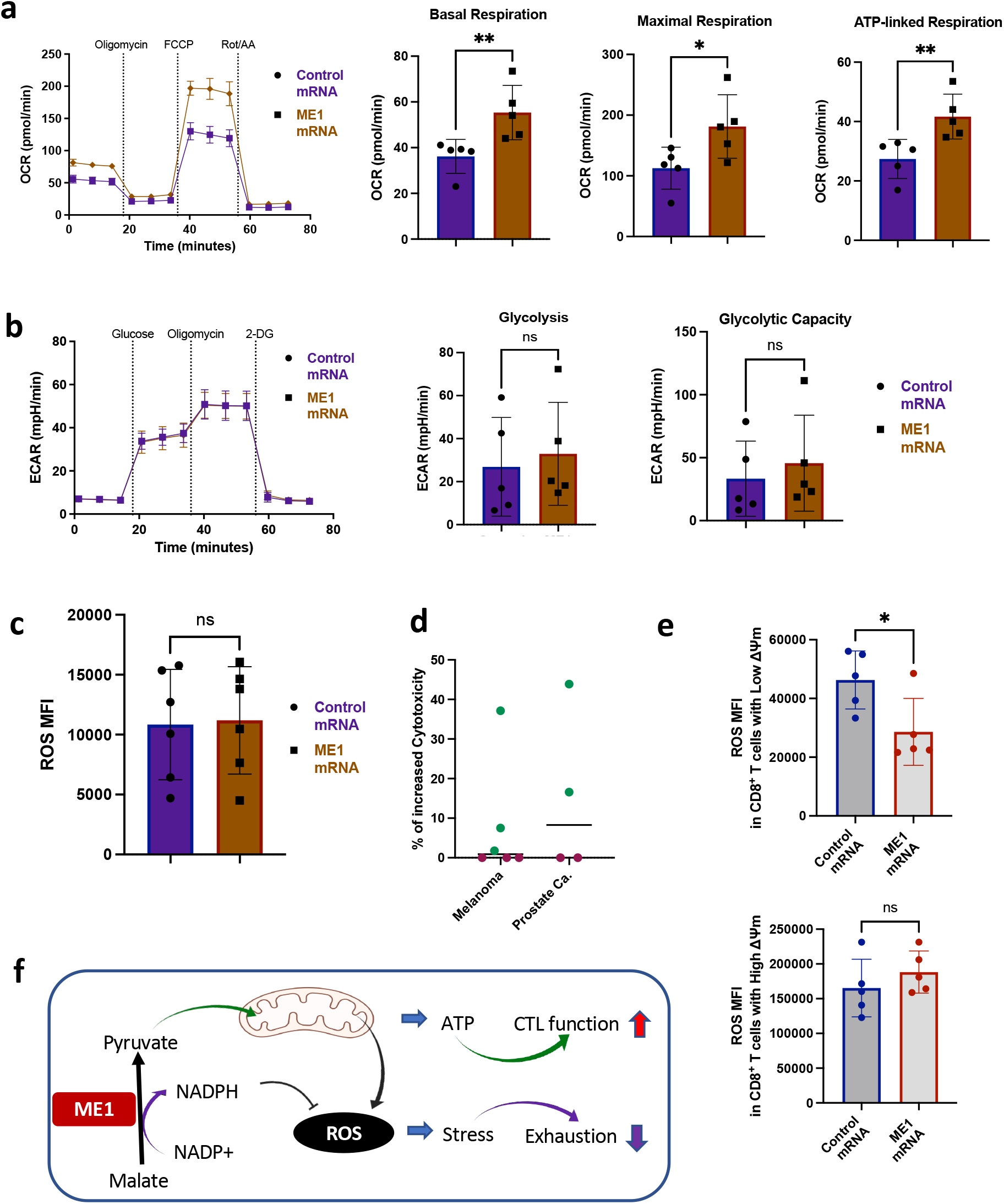
ME1 increases glycolysis and oxidative phosphorylation, but not ROS, in CD8 T cells. **a-b**, Seahorse analysis of mitochondrial respiration and ATP production (a) and glycolysis (b) in CD8^+^ T cells following transfection with control or ME1 mRNA. **c**, Cytosolic **reactive oxygen species (**ROS) in CD8^+^ T cells following transfection with control or ME1 mRNA as measured by flow cytometry (M (median)FI). **d**, Percent of increased cytotoxicity of PBMCs transfection with ME1 mRNA over PBMCs transfected with control mRNA to normalize different basal level of cytotoxicity of each patient. Cytotoxicity assay was performed at a ratio of 1:10 of tumor:effector cells. **e**, ROS levels (MFI) measured in CD8^+^ T cells with low or high ΔΨm among PBMCs of patients (n=5) with advanced prostate cancer after transfection with control or ME1 mRNA followed with T cell activation by anti-CD3 antibody for 48 hours. **f**, Graphical illustration of the potential role of ME1 in resilient CD8^+^ T cells. Statistical significance was determined by Student’s Paired (**a, b**) or two-tailed t-test; non-parametric Mann-Whitney correction (**d, e**); *P<0.05, **P<0.01, ns: not significant. CTL = cytotoxic T lymphocytes.

To test whether ME1 may have a therapeutic potential in improving CTL function of PBMCs in patients with large tumor burdens, we introduced ME1 mRNA into PBMCs isolated from patients with advanced melanoma and prostate cancer. We found that ME1 overexpression increased the tumor killing ability of PBMCs in about 50% of patients with either cancer (**Fig. 7d**). From there, we found that ME1 overexpression reduced ROS only in CD8^+^ T cells with low ΔΨm but not in high ΔΨm cells (**Fig. 7e**), suggesting CD8^+^ T cells with low ΔΨm in patients with large tumor burdens could be more sensitive to regulation mediated by ME1 overexpression. Taken together, our results imply that ME1 can promote CTL function in activated CD8+ T cells via increased cellular ATP production and mitochondrial activity, allowing energy requirements to be met, while concurrently curtailing excessive ROS, thereby preventing exhaustion (**Fig. 7f**).

## Discussion

In this study, we characterized the features of resilient T cells in patients with advanced malignancies. We found CX3CR1^+^ CD8^+^ T cells with low ΔΨm to be endowed with functional resiliency and metabolic fitness that allows them to withstand large tumor burden, retain cytotoxic functionality, and be responsive to radiation therapy or ICI treatment. One potential underlying mechanism of this resiliency is the upregulation of ME1 that not only balances excessive ROS in activated T cells, but also sufficiently enhances cytotoxic T cell responses. Thus, our study reveals a mechanism by which a highly cytotoxic function can be maintained in resilient T cells in patients with large tumor burdens and provides a method to improve the efficacy of cancer immunotherapy and radiation therapy for patients with advanced cancer.

Since PD-1 signals dynamically influence mitochondrial functions resulting in compromised function of effector T cells along with large tumor burdens^17–22^, we measured the mitochondrial membrane potential (ΔΨm) of CX3CR1^+^ CD8^+^ T cells to determine the functional state of their mitochondria, as a start to understanding T cell resilience. According to the level of ΔΨm, we found CX3CR1^+^ CD8^+^ T cells with low ΔΨm to be more responsive to radiation therapy in patients with large tumor burdens. From there, we further characterized CD8^+^ T cells with low ΔΨm and found that they are highly cytotoxic and produce less ROS. We found that ME1 is upregulated in CD8^+^ T cells with low ΔΨm and is responsible for balancing elevated levels of ROS in activated T cells and consequently promoting tumoricidal activity of CD8^+^ T cells in both healthy donors and patients with advanced cancers. To our knowledge, the role of ME1 in T cell function has not been clearly defined, although ME1 has been identified as a key tumorigenic factor by promoting the survival of tumor cells in their microenvironment where high ROS is prevalent ^39,43^. ME1 is an NADP+ dependent cytosolic enzyme that is required for catalytic activity of a reaction that converts malate into pyruvate with NAPDH as a by-product^39,40^. This means that the catalytic function of ME1 in CD8^+^ T cells with low ΔΨm may serve to provide pyruvate, which feeds into either glycolytic or oxidative phosphorylation pathways for ATP production, and thus providing an optimal maintenance of metabolic fitness in T cells. Additionally, NADPH produced from the ME-1-mediated reaction can act as an antioxidant to prevent accumulation of excessive ROS, which may trigger T cell death ^44^ and T cell exhaustion ^22,37,38^. Of note, although ME1 has the potential to reduce cytosolic ROS, probably through NADPH, our data suggest ME1 is not a scavenger of cytosolic or mitochondrial ROS in T cells that require a basal level of ROS for activation^41^.

Large tumor burden has been significantly correlated with more T cells exhaustion and poorer response to ICI therapy ^45,46^. However, the presence of resilient T cells that can recover their CTL function quickly after radiation therapy is an unexpected discovery in our study. Although the concept of resilient T cells in response to cancer immunotherapy has been conceived recently ^5–7^, no markers have been defined to identify them in patients with advanced diseases. To that end, our study suggest that CX3CR1^+^ CD8^+^ T cells with low ΔΨm could be a surrogate of resilient T cells, based on several features we identified in this report, such as (1) the quick recovery of their CTL function (degranulation/CD107a expression) after reduction of tumor burden by radiation therapy; (2) the maintenance of a low glycolytic profile, but production of optimal ATP levels; (3) the avoidance of exhaustion due to lower expression of PD-1 and TOX; (4) the shared feature of stem-like T cells (TCF1^+^PD-1^+^) which can be responsive to ICI therapy^47–52^, which is consistent with murine CD8^+^ T cells with low ΔΨm ^23^; (5) the maintenance of a relatively lower ROS level, to avoid exhaustion while not compromising T cell activation^41^; (6) the upregulation of ME1 expression, effectively balancing excessive ROS and contributing to expansion of highly cytotoxic effector T cells. These unique features of resilient T cells can not only be used as a T cell biomarker to predict or monitor patient responses to cancer immunotherapy or combination therapy, but also can be used to convey a resilient functional state of T cells in patients who may be lacking enough resilient T cells required for immunotherapeutic response. To that end, our study suggests that introduction of ME1 mRNA into T cells could be used to improve the efficacy of cancer immunotherapy either in response to ICI therapy or in adoptive T cell therapy.

Several limitations in our studies should be addressed in future studies. Although our current studies are mainly focused on the circulating CD8^+^ T cells in patients with advanced diseases, it is warranted to investigate whether tumoral CD8^+^ T cells share the features of resilient T cells. On the other hand, since there are no animal models to address the T cell-specific role of ME1 in antitumor immunity, new animal models of ME1 are needed to test how a persistent or transient expression of ME1 in tumor-reactive T cells would contribute to either endogenous tumor control or to response of ICI therapy. Also, the efficacy and safety of transferring ME1-modified T cells in treatment of established tumors would still need to be addressed by *in vivo* murine tumor experiments. Finally, one open question remains as to why some CD8^+^ T cells are endowed with or maintain a low or high ΔΨm status. It is not clear to what degree the heterogeneity of ΔΨm within human circulating CD8^+^ T cells would be related to systemic antitumor immunity, as resilient T cells need to maintain a depolarized low ΔΨm to avoid the consequences associated with high ΔΨm, such as increased ROS or exhaustion. By adapting to this unique feature, resilient T cells can protect the organism from metastatic cancer and chronic viral infections.

In summary, our study identified a mechanism by which resilient T cells can avoid exhaustion in patients with advanced cancers. Leveraging the knowledge of T cell resiliency may improve the efficacy of ICI, CAR-T cell, or TCR-T cell therapies to control metastatic malignancies that require robust systemic anti-tumor immunity for their control.

**Extended Data Fig. 1:**
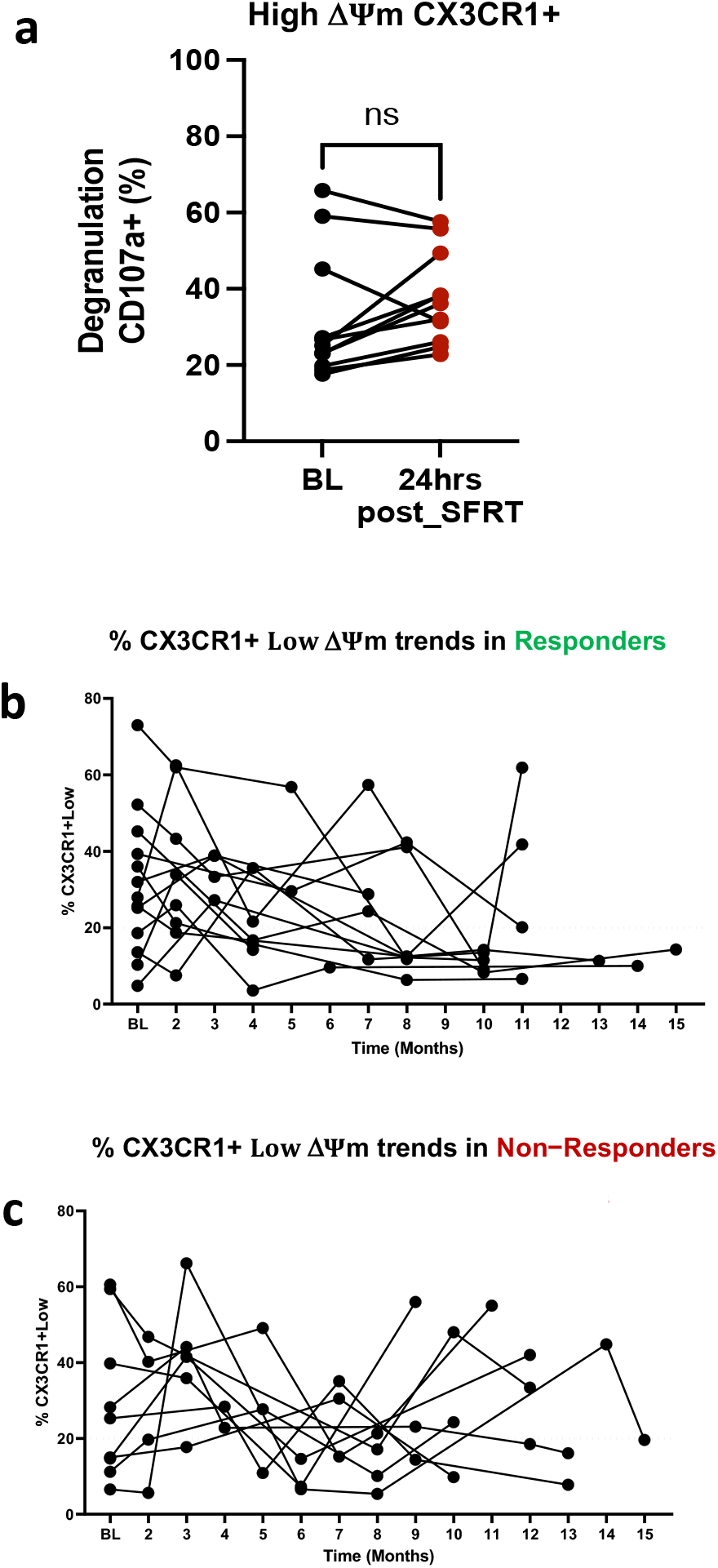
Functionality and frequency of CX3CR1^+^ CD8^+^ T cells with low or high ΔΨm in response to radiation or ICI therapy. **a**, Degranulation analysis of CX3CR1^+^ CD8^+^ T cells with high ΔΨm from patients with advanced lung cancer and sarcoma (n=17) as measured by CD107a expression comparing baseline and 24 hours (hrs) post-spatially fractionated radiotherapy (SFRT) therapy. Frequency of CX3CR1^+^ CD8^+^ T cells with low ΔΨm was measured in the peripheral blood of patients with advanced melanoma at baseline (BL) and follow up time points (N=42) in responders (**b**) and non-responders (**c**) to Immune checkpoint inhibitors (ICI) therapy. Statistical significance was determined by Student’s Paired two-tailed t-test in **a** *P<0.05, ns: not significant.

**Extended Data Fig. 2:**
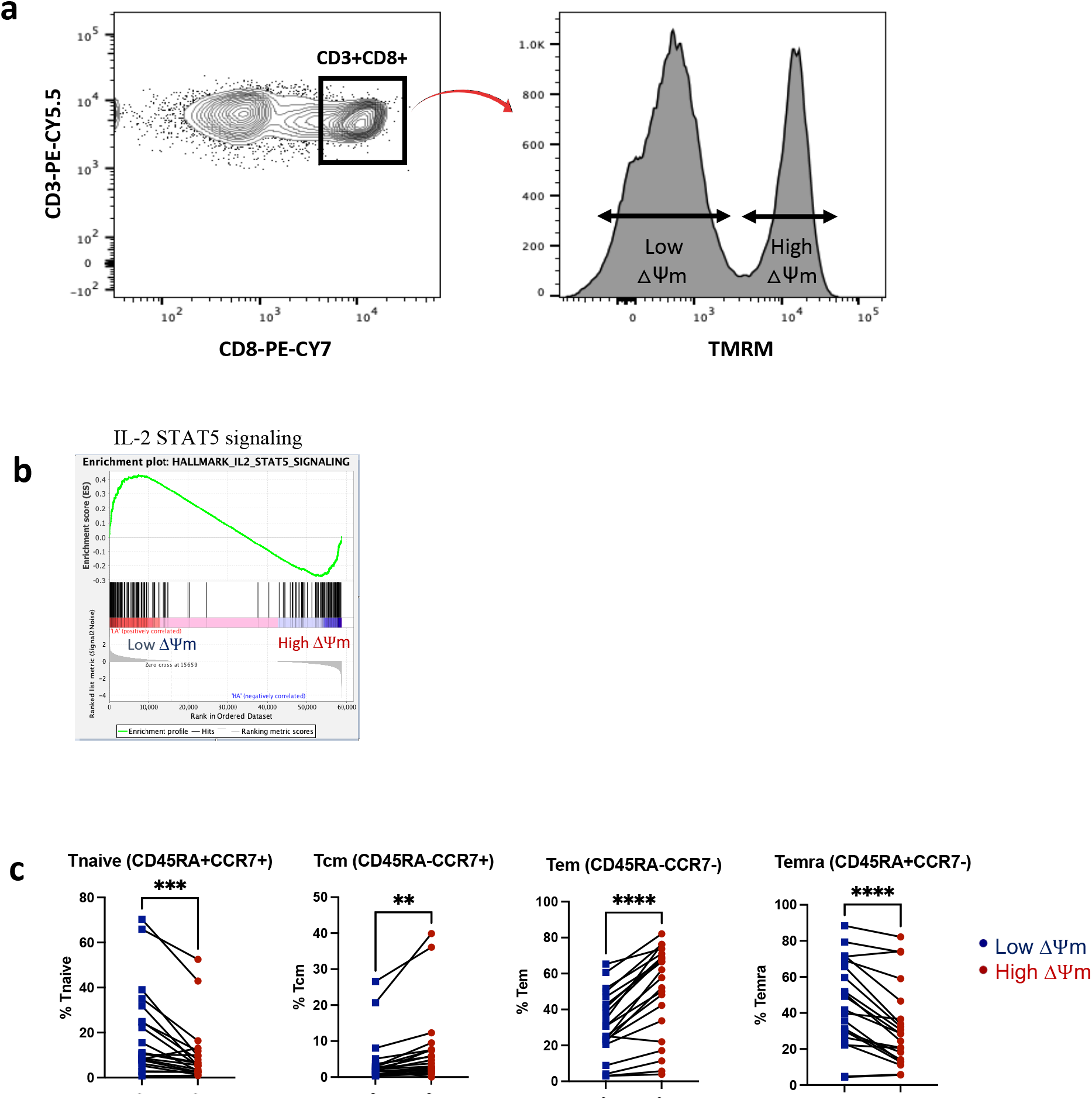
Sorting and phenotype analysis of CD8^+^ T cells with low or high ΔΨm. **a**, Gating strategy that defines the low and high ΔΨm among CD8^+^ T cells for sorting and analysis. **b**, Hallmark Gene Set Enrichment Analysis (GSEA) of RNA-sequencing data from three healthy donors using IPA analysis software. **c**, Phenotype of CD8^+^ T cell subsets as defined by CCR7 and CD45RA expression among low or high ΔΨm cells. Statistical significance was determined by Student’s Paired two-tailed t-test in **c**, **P<0.01, ***P<0.001, ****P<0.0001.

**Extended Data Fig. 3:**
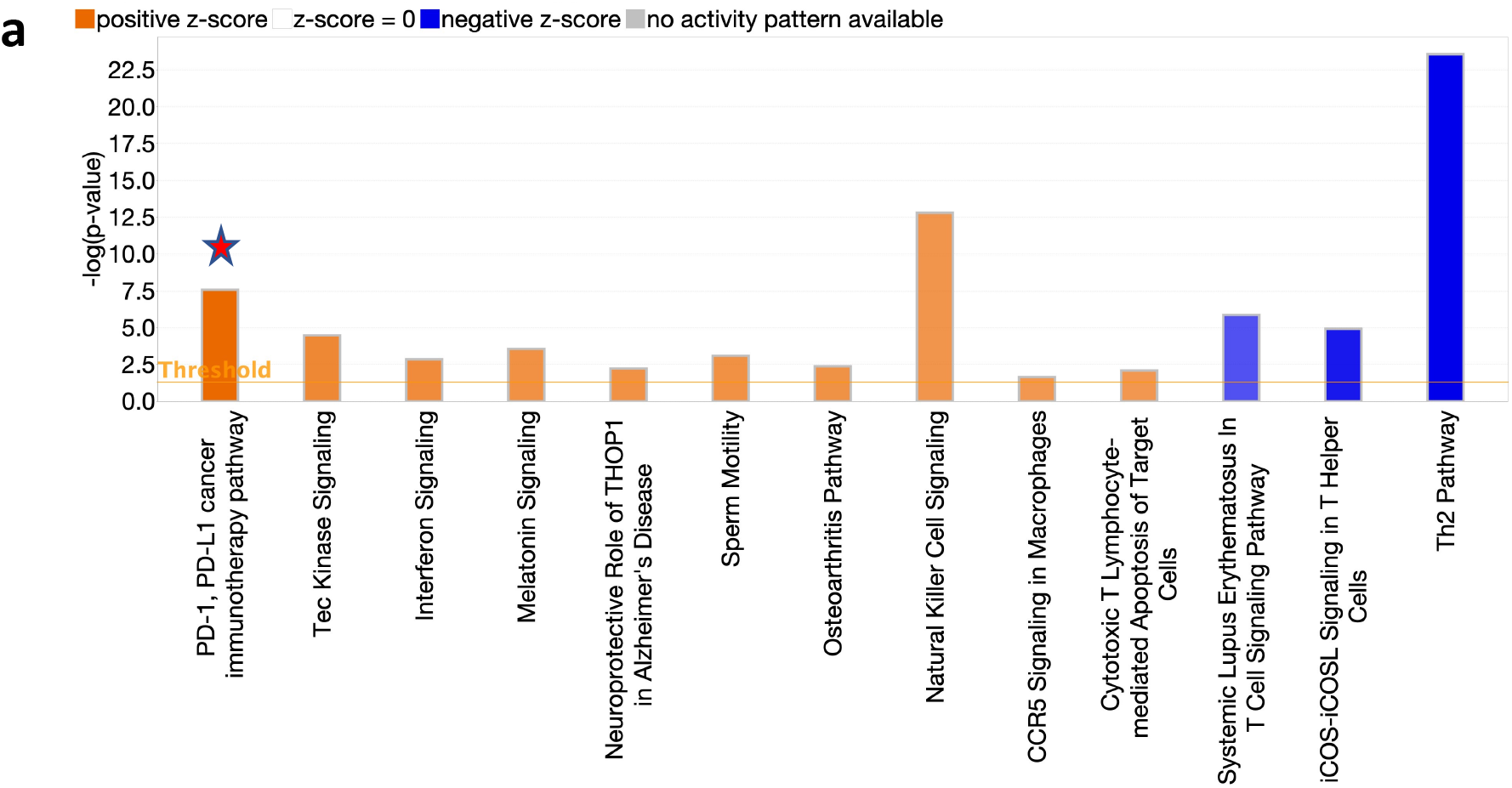
PD-1/L1 pathway is upregulated in CD8^+^ T cells with low ΔΨm cells. Hallmark Gene Set Enrichment Analysis (GSEA) of RNA-seq data from three healthy donors was further analyzed with IPA analysis software for pathway regulation. Star indicates the upregulation of PD-1/PD-L1 pathway in low ΔΨm cells.

**Extended Data Fig. 4:**
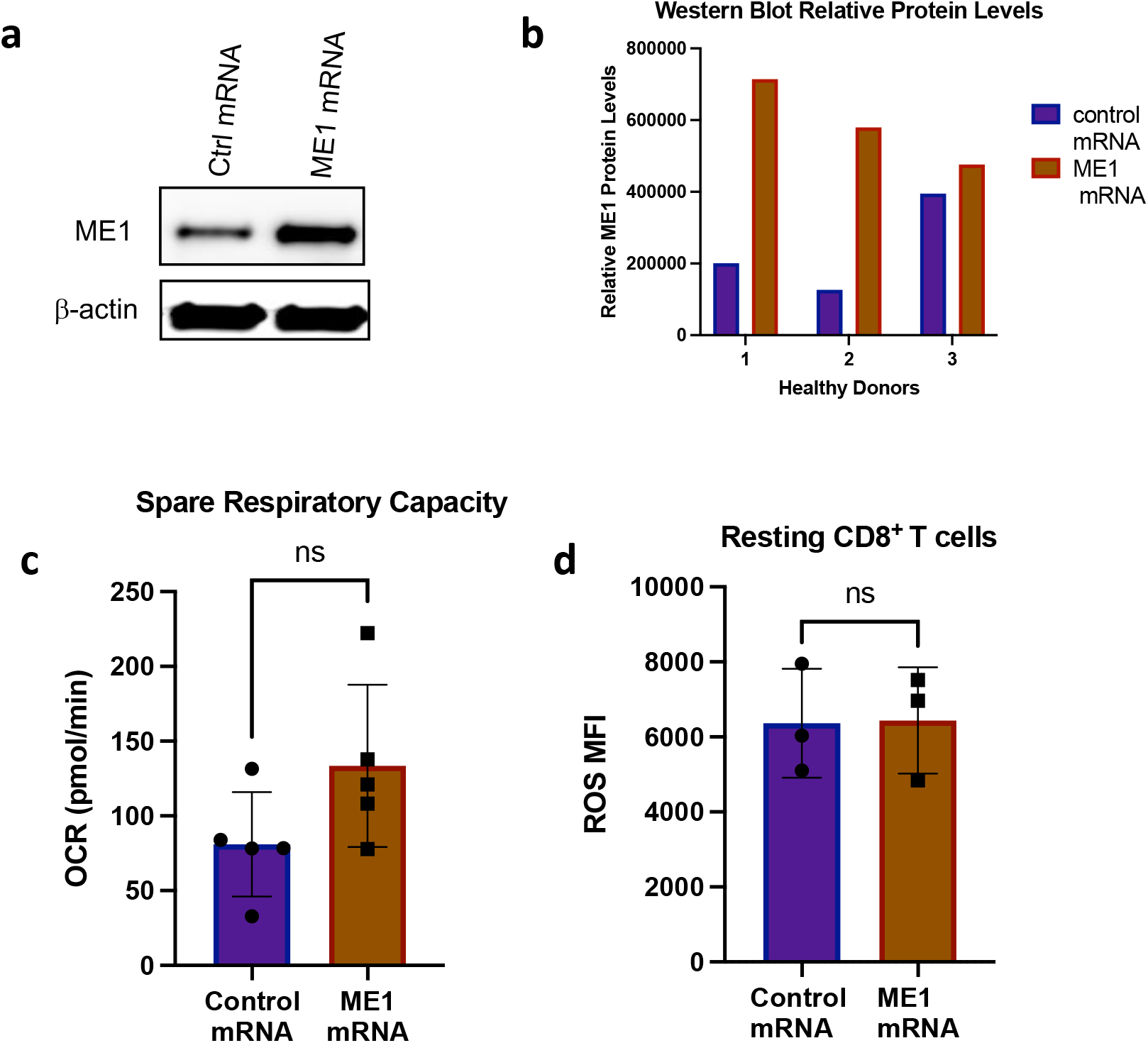
ME1 expression and its regulation of metabolism in CD8^+^ T cells. **a-b**, ME1 overexpression measured by Western blot (**a**) followed with quantification of ME1 proteins (**b**) in CD8^+^ T cells after transfection with control or ME1 mRNA. **c**, Spare respiratory capacity as measured via Seahorse assay. **d**, Reactive oxygen species (ROS) M (median)FI in resting CD8^+^ T cells overexpressing ME1 versus control mRNA. Statistical significance was determined by Student’s unpaired (**c, d**) two-tailed t-test; non-parametric Mann-Whitney correction, ns: not significant.

## METHODS

### Specimen collection and T cells isolation

Peripheral blood mononuclear cells (PBMCs) were isolated from healthy donors or patients via centrifugation with lymphoprep (07851, StemCell Technologies) and SepMate conical tubes (85450, StemCell Technologies). CD8^+^ T cells were then isolated using a magnet based CD8 T cell isolation kit (19053, StemCell Technologies) and used immediately for experiments. Some experiments used PBMCs (patient samples) previously stored in liquid nitrogen. After thawing, PBMCs would be incubated in CTL medium (RPMI 1640 complete medium, rhIL-2: 10 U/ml, rh IL-15: 5 ng/ml; rhIL-7: 5 ng/ml) at 37°C overnight for recovery before transfection.

#### Patient information

Peripheral blood for this study were collected after written consent was obtained from each participant. Clinical course, treatment information, and outcomes in patients treated with anti-PD-1/L1 therapy and radiation therapy at Mayo Clinic were retrospectively collected. Response to treatment was evaluated according to standard clinical practice guidelines using RECIST ^11^. Note that the response listed for anti-PD-1/PD-L1 therapy, (R = complete response, NR = progressive disease, SD = stable disease) was apparent (using RECIST) at the 12-week post-initiation of therapy time-point for all patients. For prostate cancer (receiving SFRT and SBRT), the response was evaluated based on the PSA levels and distance reoccurrence.

#### Study approval

Peripheral blood for this study from healthy people were acquired from anonymous donors who had consented for blood donation at the Blood Transfusion Center at Mayo Clinic. All donors and patients provided signed informed written consent; the study was approved by the Mayo Clinic Rochester IRB and was conducted according to Declaration of Helsinki principles.

### TMRM Staining and Cell Sorting

CD8^+^ T cells were washed once with 1x PBS and adjusted to a concentration of 1 × 10^6^ cells/ml and stained with 0.02 μM final concentration of tetramethylrhodamine methyl ester (TMRM) or 2 nM final concentration of Carbonyl Cyanide 3-chlorophenylhydrazone (CCCP) as control. The TMRM stained cells were incubated at 37°C for 30 minutes with intermittent shaking while the CCCP stained cells were incubated at 37°C for 5 minutes. The cells were then washed twice with 1x PBS and resuspended at 10-15 × 10^6^ cells/ml of cell culture media for sorting. The cells were sorted with the BD FACSMelody^TM^ Cell Sorter using the 100-micron sort nozzle and on the PE channel (561nm excitation).

### siRNA Transfection

Sorted CD8+ T cells with low and high ΔYm were centrifuged at 200 × g for 10 minutes prior to nucleofection (4D nucleofector system, Lonza). Three-five million cells were combined with 200 pMol siRNA (siControl or siME1) in 20 μl P3 nucleofection media (Lonza) per well of the 16-well Nucleocuvette strips (X unit). Program FI-115 was used. Following nucleofection, cells were rested in warm RPMI (no FBS, no cytokines) for 4 hours before adding 10% FBS and 10 IU/ml of IL-2, 5 ng/ml of IL-7, and 5 ng/ml of IL-15 into the culture for an overnight recovery followed with subsequent inclusion in experiments.

### mRNA Transfection

T cells were transfected with control mRNA or ME1 mRNA (220 μg/ml) using the P3 nucleofection kit (Lonza V4XP-3024) and program FI-115 on the 4D Nucleofector (Lonza). Following nucleofection, cells were rested in warm RPMI (no FBS, no cytokines) for 4 hours before adding 2x CTL medium (as described above) for an overnight recovery followed with T cell activation and functional analysis.

### Cytotoxicity Assay

Target tumor cells were washed twice with HBSS, and then labeled with Calcein-AM (5 μM) for 30 minutes and incubation at 37°C in dark, shaking occasionally. The cells were then again washed with HBSS twice and re-suspended at 1 × 10^5^/ml in CTL medium with no FBS. Preactivated CD8^+^ T cells and target cells were mixed at 1:20 or 1:10 (target to effector ratio) in CTL media without FBS and seeded into 96-U bottom well plate at 200 μl per well. Saponin (0.1%) or Triton X-100 (2%) was added to wells that contained only tumor cells to provide a value for “maximum calcein release,” in each assay; tumor cell-only wells were included to measure spontaneous release of calcein. All experimental and control conditions were performed in triplicate wells. The plate was briefly centrifuged at 1,000 rpm for 30 seconds, followed by incubation at 37°C for 4 hours. The plate was then centrifuged at 2,000 rpm for 5 minutes. 100 μl of the 200 μl supernatant was then removed from each well and added to a new, opaque (black) 96 well flat bottom plate (Thermo scientific, 237105). Calcein fluorescence was read using an automated fluorescence measurement system (BioTeK Synergy HTX multi-mode reader) with an excitation of 485/20 and an emission filter of 530/25 scanning for 1 second per well. Percent cytotoxicity was then calculated using the following formula:

% Cytotoxicity = 100 × [(Experimental well release-Spontaneous release well)/Maximum release well-Spontaneous release well].

### Degranulation assay

T cells were adjusted at a concentration of 1 × 10^6^ cells/100 μl CTL medium including Golgi-Stop (Biolegend, 420701) and Golgi-Plug (Biolegend, 420601) and incubated with 5 μl of CD107a antibody (Biolegend, H4A3) and 5 μl of anti-CD3/CD28 beads. The cells were then briefly centrifuged at 300 × g for 1 minute and incubated at 37°C for 5 hours. After the incubation, the cells were stained with TMRM followed with surface antibody staining before flow cytometry analysis.

### ROS detection

Cells were stained with 250 nM of CellROX (ThermoFisher Scientific, C10492 or 1 μM of MitoSOX (ThermoFisher Scientific, M36008) in complete media and incubated at 37°C for 45 minutes. From there, the cells were resuspended in FACS buffer (1x PBS, 2mM EDTA, and 3% FBS) at a concentration of 1 × 10^6^ cells/ 100 μl followed by staining with antibodies for surface molecules for 20 minutes at room temperature in the dark. Cells were then washed once in FACS buffer, resuspended in 200 μl, and analyzed on Bio-Rad ZE5 Cell Analyzer. Analysis was performed using FlowJo V10.

### Flow Cytometry Analysis

Cells were adjusted to 0.5-1 × 10^6^ cells/ml with 1x PBS and stained with live/dead dye and incubated at 4°C for 30 minutes. The cells were then washed once with 1x PBS followed by staining for cell surface molecules. For intracellular molecule staining, cells were incubated with FoxP3 Fixation Buffer overnight at 4°C. After fixation, cells were washed once with 1x permeabilization buffer, and then stained with antibodies for intracellular molecules such as ME1, TCF-1/7, TOX, Eomes, T-bet, Ki67, HIF1-α, NKG7, and Glut1 for 1 hour at 4°C in 100 μl of 1x permeabilization buffer. Following intracellular staining, cells were washed once with 1x permeabilization buffer and then resuspended in 200 μl FACS buffer for flow cytometry analysis, which was completed on a Cytoflex LX (Beckman Coulter) (DAQ Version V2.233, MCB Version: V3.01) running CytExpert software. Flow cytometric analysis was performed using FlowJo V10.

### Quantitative PCR

RNA was isolated using Qiagen RNeasy Plus Mini Kit (74134, Qiagen). RT-reaction was completed using SuperScript™ III Reverse Transcriptase (18080-400, Invitrogen). qRT-PCR analysis was completed using a Quant-Studio 3 Real-Time PCR System (Applied Biosystems) and TaqMan Fast Advanced Master Mix (444455, Applied Biosystems). Taqman assays included: Hs01120688-gl, Human ME1-FAM and Hs99999901-Sl, Human-18s-FAM, both from Applied Biosystems. All samples were run in doublet. Relative expression was calculated using the 2^-ΔCt^ method. Granzyme B (F 5-TACCATTGAGTTGTGCGTGGG-3, R 5-GCCATTGTTTCGTCCATAGGAGA-3, ME1 (F 5-GGGAGACCTTGGCTGTAATGG-3, R 5-TTCGGTTCCCACATCCAGAAT-3), UBE (F 5-GTACTCTTGTCCATCTGTTCTCTG-3; R 5-CCATTCCCGAGCTATTCTGTT-3). RNA input was normalized to 20 ng/ul with 10 ul input for the RT reaction. Post RT reaction cDNA was diluted 1:5 before amplification on the Quantstudio 3 in the following volume per well: 5 μl cDNA template, 10 μl SYBR green (Applied Biosystems), 3 μl H2O, 1 μl 10 mM Forward/Reverse primer.

### Western Blots

Cell pellets were lysed in NP-40 buffer and concentrations measured via protein assay using BioRad reagent (#500-0006). Once diluted to equal concentrations, samples were combined with Laemmli sample buffer (BioRad) with beta-mercaptoethanol and run using a Mini-PROTEAN Electrophoresis and Transfer system (BioRad). Membranes were incubated with primary antibody overnight at 4°C (ME1, Beta-Actin, or GAPDH). Secondary antibodies were added the next morning for 2 hours (HRP-conjugated anti-mouse). Signal was visualized using SuperSignal West Pico PLUS Chemiluminescence Substrate Kit (34577, Thermo Fisher) on a Syngene G: Box Chemi XX6 system running GeneSys software (V1.6.1.0). Full images of all membranes are included in the supplementary source data files.

### Transmission electron microscopy (TEM) analysis of mitochondria

Cells were fixed in Trump fixative for 1 hour at room temperature or at 4°C overnight followed with a fixation for 1 hour in 1% osmium tetroxide. The samples were dehydrated, embedded in Spurrs resin, sectioned at 90 nm, and observed using a Joel 1400 electron microscope (Joel USA Inc.). For quantification, images of individual T cell in a single field of view downloaded into JPEG images and the number of mitochondria structures within the T cells were counted manually by two different readers.

### Metabolic Assays

Seahorse Xfe96 Bioanalyzer (Agilent) was used to determine OCR and ECAR. Sorted cells or total CD8^+^ T cells were washed in XF Base media (Seahorse XF RPMI medium with 2 mM glutamine, 10 mM glucose, 1 mM sodium pyruvate, and 5 mM HEPES (4-(2-hydroxyethyl)-1-piperazineethanesulfonic acid), pH 7.4 at 37 °C) for OCR or XF media with 2 mM glutamine for ECAR before being plated onto Seahorse cell culture plates coated with Cell-Tak (Corning #354240) at 1 × 10^5^ cells per well. The cells were allowed to adhere to culture plate. The OCR was measured via Seahorse Mito Stress assay (Agilent), with addition of oligomycin (2 μM), carbonyl cyanide 4-(trifluoromethoxy) phenylhydrazone (FCCP; 1.2 μM) and Rotenone and Antimycin (1.0 μM)). The ECAR was measured with addition of 10 mM glucose, 2 μM oligomycin, and 50 nM 2-deoxy-D-glucose (2-DG). Assay parameters were as follows: 3-minute mix, no wait, 3-minute measurement, repeated 3–4 times at basal and after each addition. SRC was calculated as oxygen consumption rate (OCR) at maximum rate (OCRMax) - OCR in basal state (OCRBas). Mitochondrial ATP production was calculated by subtracting the minimum respiration rate after oligomycin injection from the basal respiration rate before oligomycin injection.

### Mitochondria mass Staining

Cells at a concentration of 1 × 10^6^ cell/ml were washed once with 1x PBS and incubated with 100 nM of MitoTracker Green FM dye for 30 minutes at 37 °C. The cells were then analyzed by flow cytometry.

### Bulk RNA sequencing

RNA was isolated using Qiagen Rneasy Plus Mini Kit (74134, Qiagen). The bulk RNA paired-end sequencing reads were processed through the Mayo bioinformatics pipelines MAP-Rseq (v3.0) as we previously reported. Reads were aligned to human reference genomes (hg38). The RNA aligned reads were quantified for gene expression using the Subread package. Differences across groups were assessed using bioinformatics package edgeR 2.6.2 to identify differentially expressed genes. Such genes were reported with magnitude of change (log2 scale) and their level of significance (False Discovery Rate, FDR < 5%). For pathway analysis, T cells with low or high ΔYm were randomized and put through Gene Set Enrichment Analysis (GSEA) as described in the user guide.

### Statistical Analyses

Data were analyzed in GraphPad Prism (version 9) using the unpaired or paired, two-tailed *t*-test without correction for multiple comparisons, as indicated in figure legends.

### Antibody List

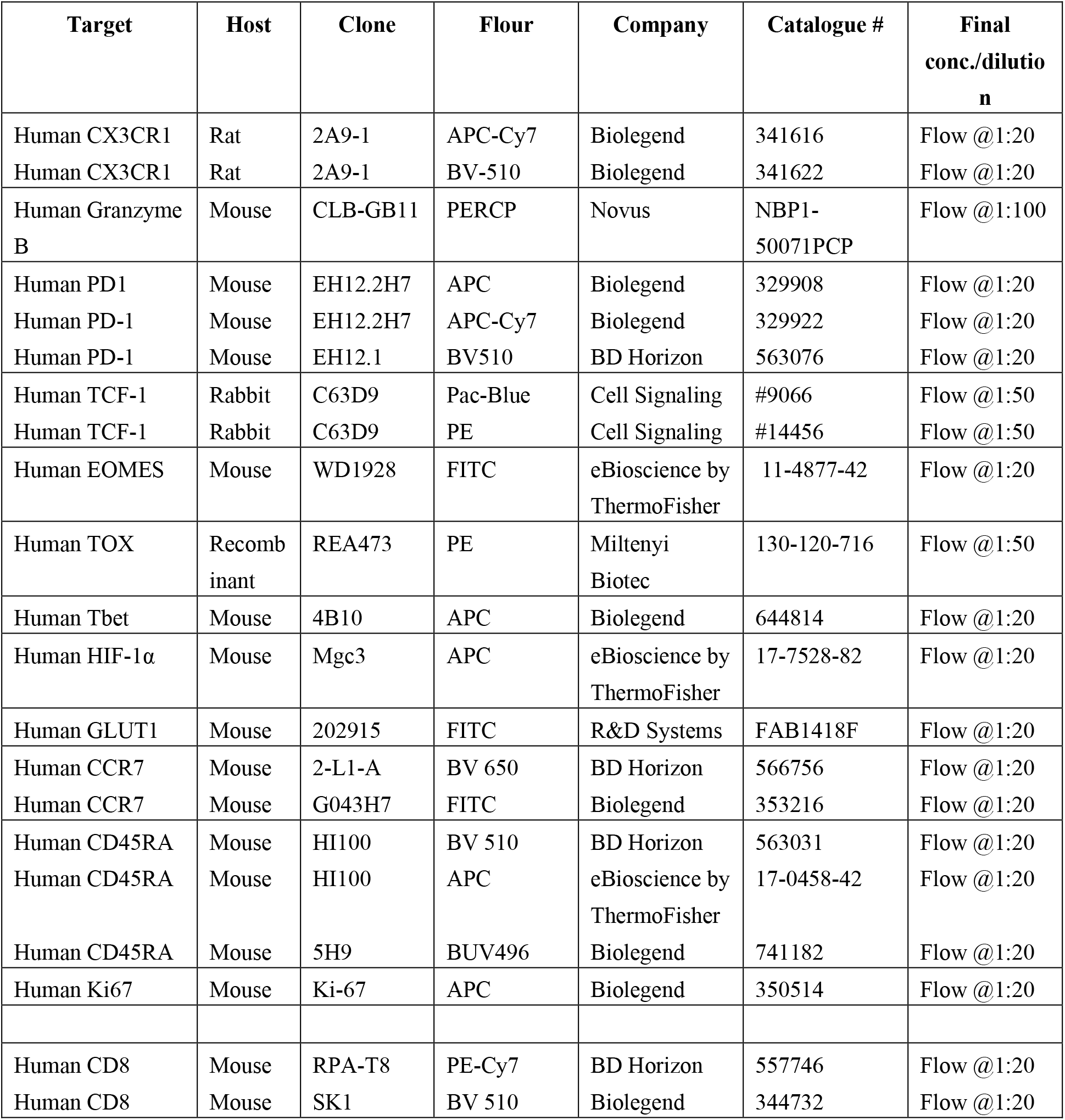
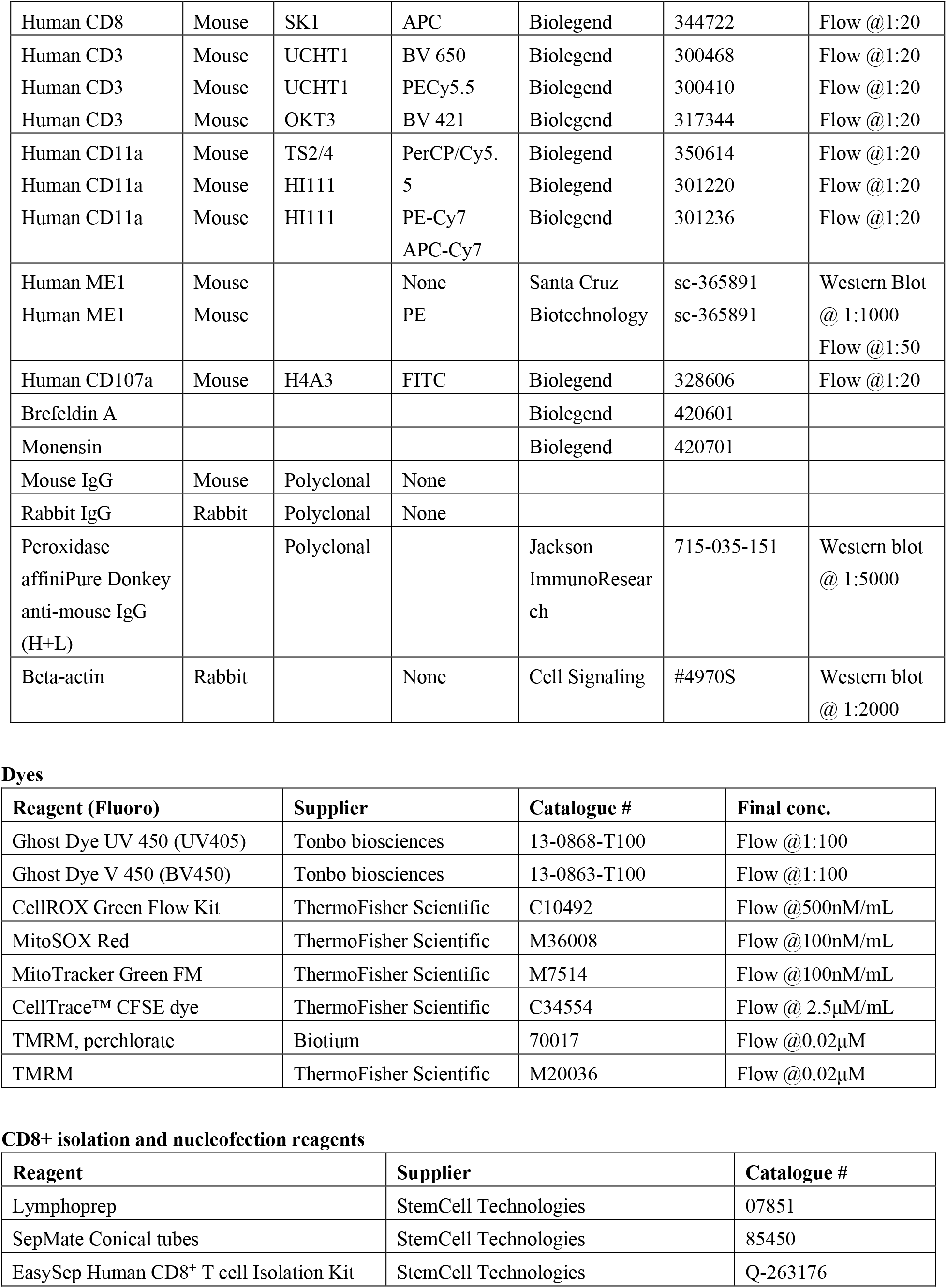
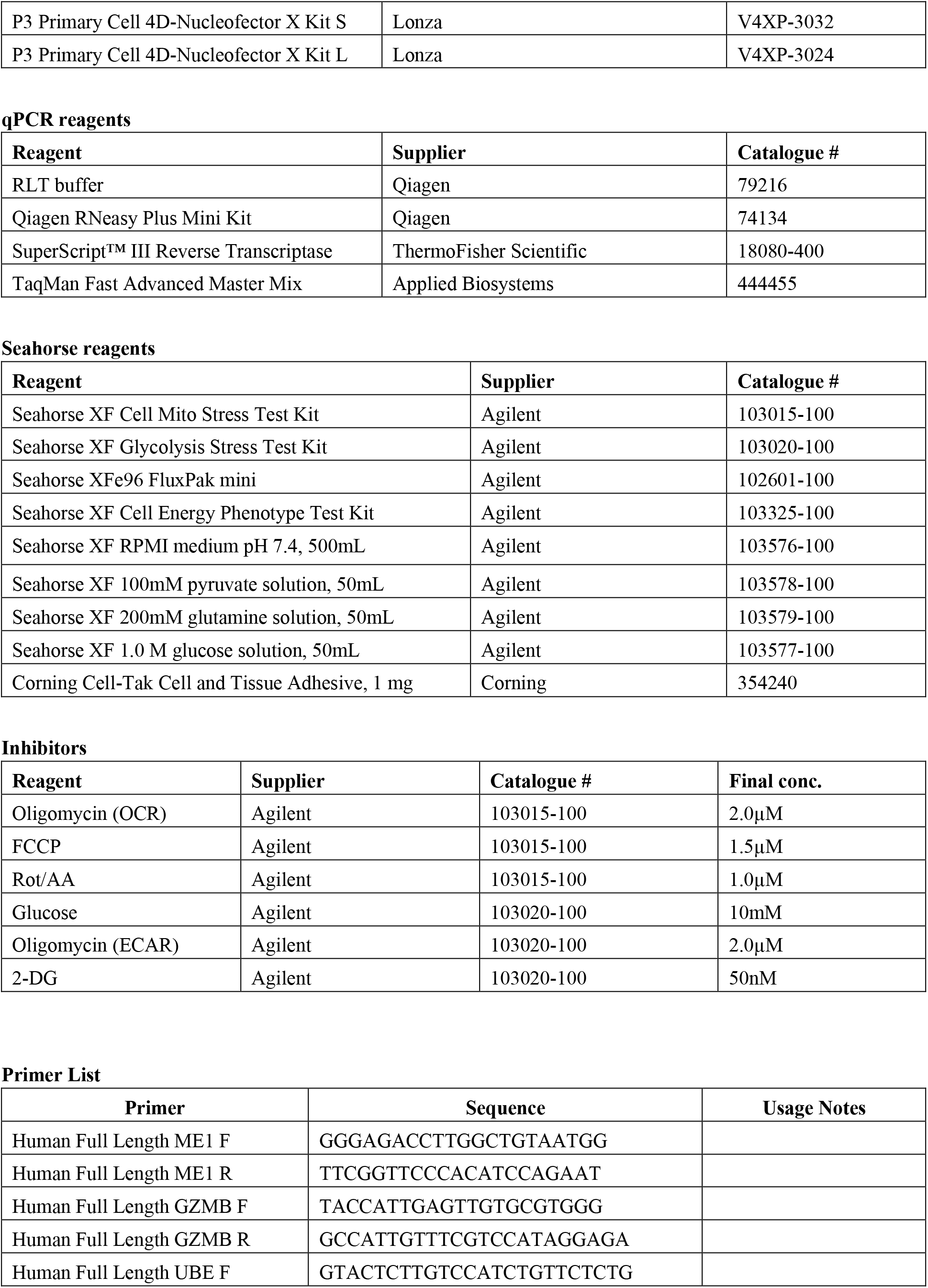
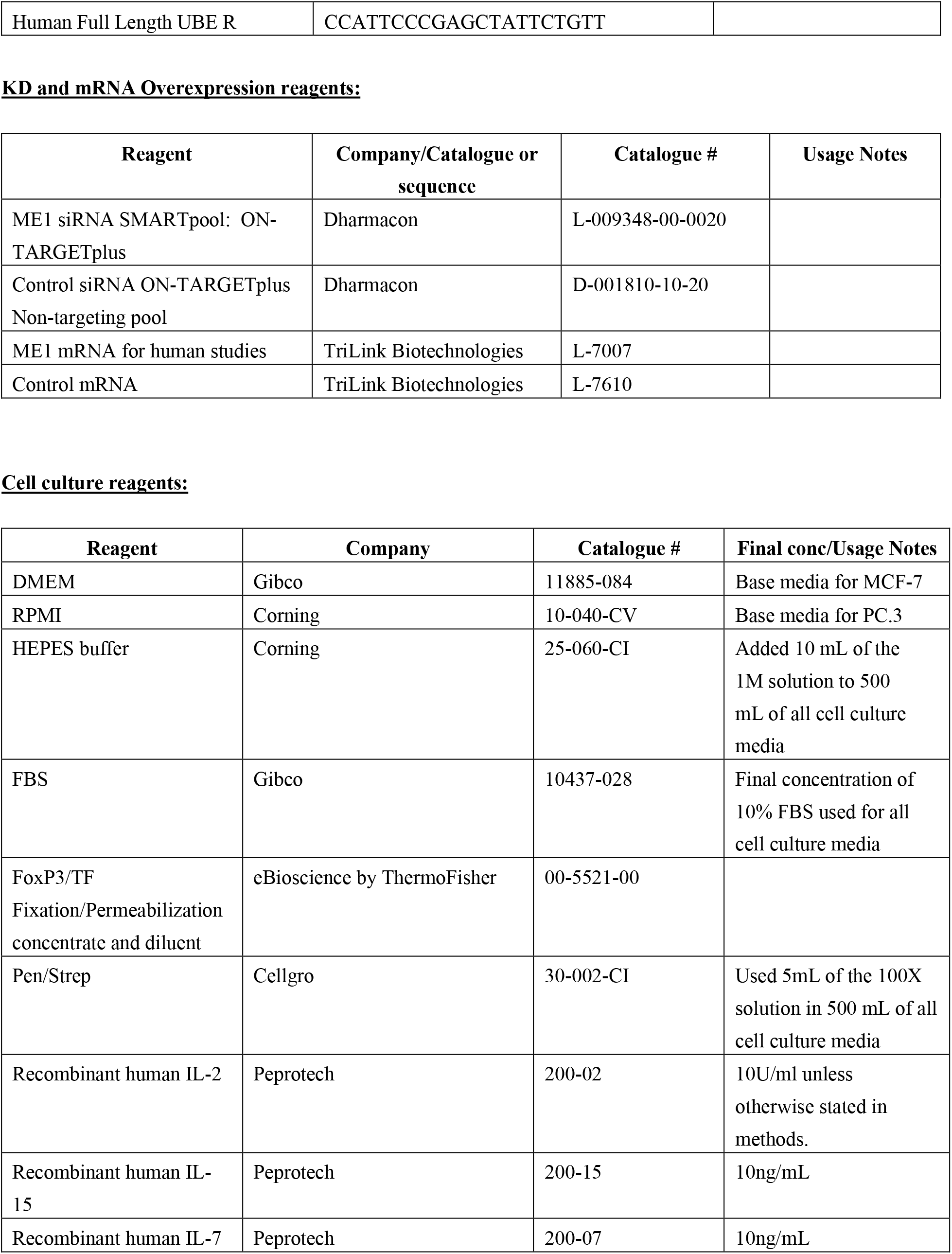
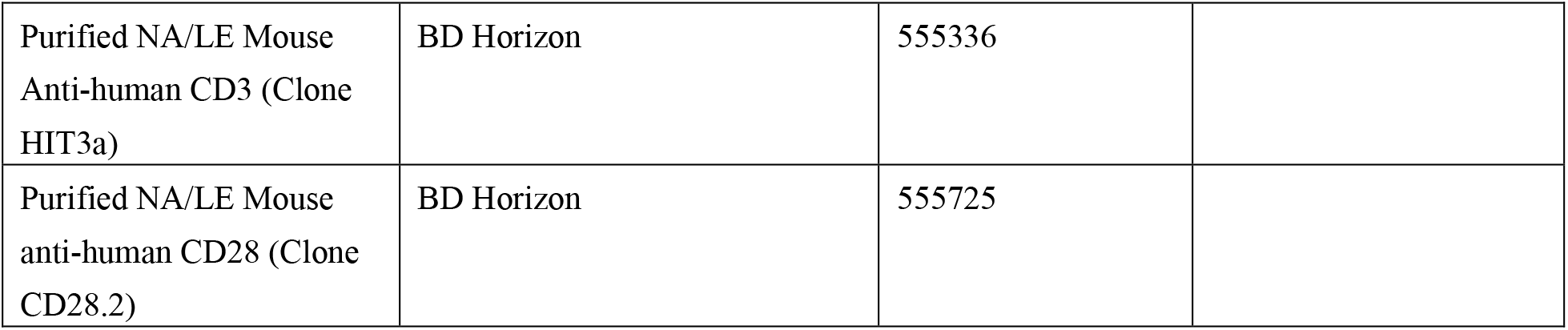

## Acknowledgements

We thank all the patients who generously contributed samples and participated in the study. We thank all members of the Dong laboratory for helpful discussions and critical analysis of the manuscript. We also acknowledge the Department of Radiation Oncology and Division of Medical Oncology at the Mayo Clinic for providing excellent patient care. This study was supported in part by the generosity of Mayo Clinic benefactors. We acknowledge the Genome Analysis Core at Mayo Clinic, including technologists Vernadette A. Simon and Fariborz Rakhshan-Rohakhtar as well as supervisors Julie S. Lau, Samantha J. McDonough, and Mark Mutawe. We are grateful for the electron microscopy imaging service provided by Bing Huang and Scott I. Gamb; and Flow cytometry and Sorting services provided by Holly Lamb, Colleen Moe, Han Yong Hwan, and Drew Kluge at the Mayo Clinic Microscope and Cell Analysis Core. The staff at Mayo Clinic Blood transfusion center. BioRender was used to create the graphics throughout the manuscript. We acknowledge support from the Mayo Clinic Center for Individualized Medicine’s IMPRESS program and High-Definition Therapeutics program (H. Dong), the Mayo Clinic Center for Biomedical Discovery (H. Dong), the Mayo Clinic Cancer Center’s David F. and Margaret T. Grohne Cancer Immunology and Immunotherapy program (H. Dong), NIH grant R21 CA251923 (A.S. Mansfield), NIH grant K12 CA 090628 (Y. Yan), NIH grant R01 AR077518 (Hu Zeng), NIH grant R21 CA197878 (H. Dong), NIH grant R01 AI095239 (H. Dong), NIH grant R01 CA200551 (H. Dong and S.S. Park), NIH grant R01 CA256927 (H. Dong), and NIH grant R01 200551 (S.S. Park and H. Dong). This project is partially funded by the Lawrence and Marilyn Matteson Award S.S. Park, D.O, and M.P. Grams), and Sydney Luckman Family Predoctoral Award (J.K.G).

## Author contributions

J.K.G. and H.D. conceived the project, designed experiments, and wrote the manuscript. J.K.G. performed most of the experiments. Z.M., C.L., J.B.H., V.V.V., E.R.D., M.A.H., W.B., and Y.H. processed patient blood samples, conducted some of the flow cytometry experiments, Western blot experiments and qRT-PCR. Y.L (Li). analyzed the bulk RNAseq data and J.K.G. ran the GSEA analysis. G.D. performed healthy donors’ ME1 overexpression cytotoxicity assays and Western blots. F.A., S.H., and J.J.O. performed statistical analysis. F.L. and H.Z. advised on biochemical and metabolic assays and provided critical edits to the manuscript. X.W. provided guidance on messenger RNA design and construct. Y.Y., R.X.D., M.Z., S.N.M., A.S.M., Y.L.(Lin)., M. G., D.O. and S.S.P. provided clinical samples and interpretation of clinical data. All authors reviewed the manuscript.

